# Systematic and functional analysis of horizontal gene transfer events in diatoms

**DOI:** 10.1101/2020.01.24.918219

**Authors:** Emmelien Vancaester, Thomas Depuydt, Cristina Maria Osuna-Cruz, Klaas Vandepoele

## Abstract

Diatoms are a diverse group of mainly photosynthetic algae, responsible for 20% of worldwide oxygen production, which can rapidly respond to favourable conditions and often outcompete other phytoplankton. We investigated the contribution of horizontal gene transfer (HGT) to its ecological success. A systematic phylogeny-based bacterial HGT detection procedure across nine sequenced diatoms showed that 3-5% of their proteome has a horizontal origin and a large influx occurred at the ancestor of diatoms. More than 90% of HGT genes are expressed, and species-specific HGT genes in *Phaeodactylum tricornutum* undergo strong purifying selection. They are implicated in several processes including environmental sensing, and expand the metabolic toolbox. Cobalamin (vitamin B12) is an essential cofactor for roughly half of the diatoms and is only produced by bacteria. Genes involved in its final synthesis were detected as HGT, including five consecutive enzymes in *Fragilariopsis cylindrus*. This might give diatoms originating from the Southern Ocean, a region typically depleted in cobalamin, a competitive advantage. Overall, we show that HGT is a prevalent mechanism that is actively used in diatoms to expand its adaptive capabilities.

## 2. Introduction

Horizontal, also dubbed lateral, gene transfer (HGT) is the transfer of genetic information between reproductively isolated species by a route other than direct exchange from parent to progeny. Although HGT events are widespread and well documented among prokaryotes, they are much rarer in eukaryotes. Nevertheless, recently several examples of HGT from archaea or bacteria into eukaryotes have been reported. Functional HGT events have been described for almost all unicellular eukaryotic lineages, including fungi^1,2^, extremophilic red algae^3^, green algae^4^, rumen-associated ciliates^5^, oomycetes^6^ and photosynthetic diatoms^7,8^. Next to events involving the maintenance of pre-existing functions, which occur mainly in endosymbiotic relationships, innovative events have been described which provide the recipient with new functions or an altered phenotype^9^. Although the uptake of genetic material happens by chance, fixation does not, making HGT predominantly important in the following processes: i) the alteration of iron uptake and metabolism^8,10,11^, ii) adaptation to an anaerobic lifestyle^12,13^, iii) nucleotide import and synthesis^1,14^, iv) novel defence mechanisms^15,16^, v) mechanisms to cope with stressors such as salt^17,18^, temperature^4^ and heavy-metal concentrations^3^ and vi) expansion of its metabolic capacities^2,5,6^.

Diatoms (Bacillariophyta) are one of the most abundant and species-rich groups of phytoplankton and release between 20-25% of the global amount of oxygen^19^. They can rapidly adapt to local conditions, outcompete other photosynthetic eukaryotes and dominate oceanic spring blooms, as long as silicon is not limited^20^. Moreover, they are found throughout every aquatic photic zone of this planet, such as oceans, intertidal zones, freshwater bodies, soil and even ice ecosystems^21^. Molecular clock evidence suggests that diatoms emerged between 225 and 200 million years ago^22^ and their origin may be related to the end-Permian mass extinction which occurred around 250 million years ago. In the early Cretaceous, between 150 and 130 million years ago, diatoms split into the centric and pennate lineage. Several whole-genome sequences of representatives from polar centrics (*Thalassiosira pseudonana*^23^, *Thalassiosira oceanica*^24^, *Cyclotella cryptica*^25^), araphid pennates (*Synedra acus*^26^) and raphid pennates (*Phaeodactylum tricornutum*^7,27^, *Fistulifera solaris*^28^, *Fragilariopsis cylindrus*^29^, *Pseudo-nitzschia multistriata*^30^) have become available in recent years, which allows the analysis of the evolutionary history within diatoms. It is not fully understood how HGT has contributed to the ecological success of this environmentally important group of organisms. Moreover, diatoms evolved from several endosymbiotic events and their plastid is thought to have originated from a red alga, which has also contributed to their genetic set-up.

Although HGT detection has been previously performed in diatoms within the context of genome projects^7,24,25,27,30^, they were based on different methodologies and criteria and are therefore not directly comparable. While some studies used phylogenetics^7,30^, others relied purely on sequence homology searches^24,25,27^. In this study, we sought to systematically detect HGT events simultaneously across all sequenced diatoms. We delineated genes from horizontal descent using a high-throughput gene family phylogenetics-based approach, which allows dating transfer events. Here, we comprehensively explore the functional bias of HGT genes in diatoms and for the first time gain insight into their expression dynamics and patterns of selection.

## 3. Results

### Detection and phylogenetic distribution of diatom HGT candidates

Twenty unicellular eukaryotic species (Table S1) were selected to deduce the contribution of bacterial-derived HGT. The identification of bacterial-to-eukaryotic HGT was achieved by building a phylogenetic tree per gene family. Therefore, all protein-coding genes from 20 eukaryotic species (Figure 1a) were clustered in 145,601 gene families, of which 32% are genes lacking similarity to any other protein in this dataset, followed by phylogenetic tree construction for 8,476 gene families having similarity to bacterial proteins. Also the species topology of these 20 unicellular eukaryotes was constructed, both based on single-locus trees and a concatenation-based approach of 156 near-single copy gene families (138,948 amino acids) (Figure 1a), with the haptophyte *Emiliania huxleyi* as an outgroup. Having the species tree available, allows for the dating of HGT candidates.

**Figure 1:**
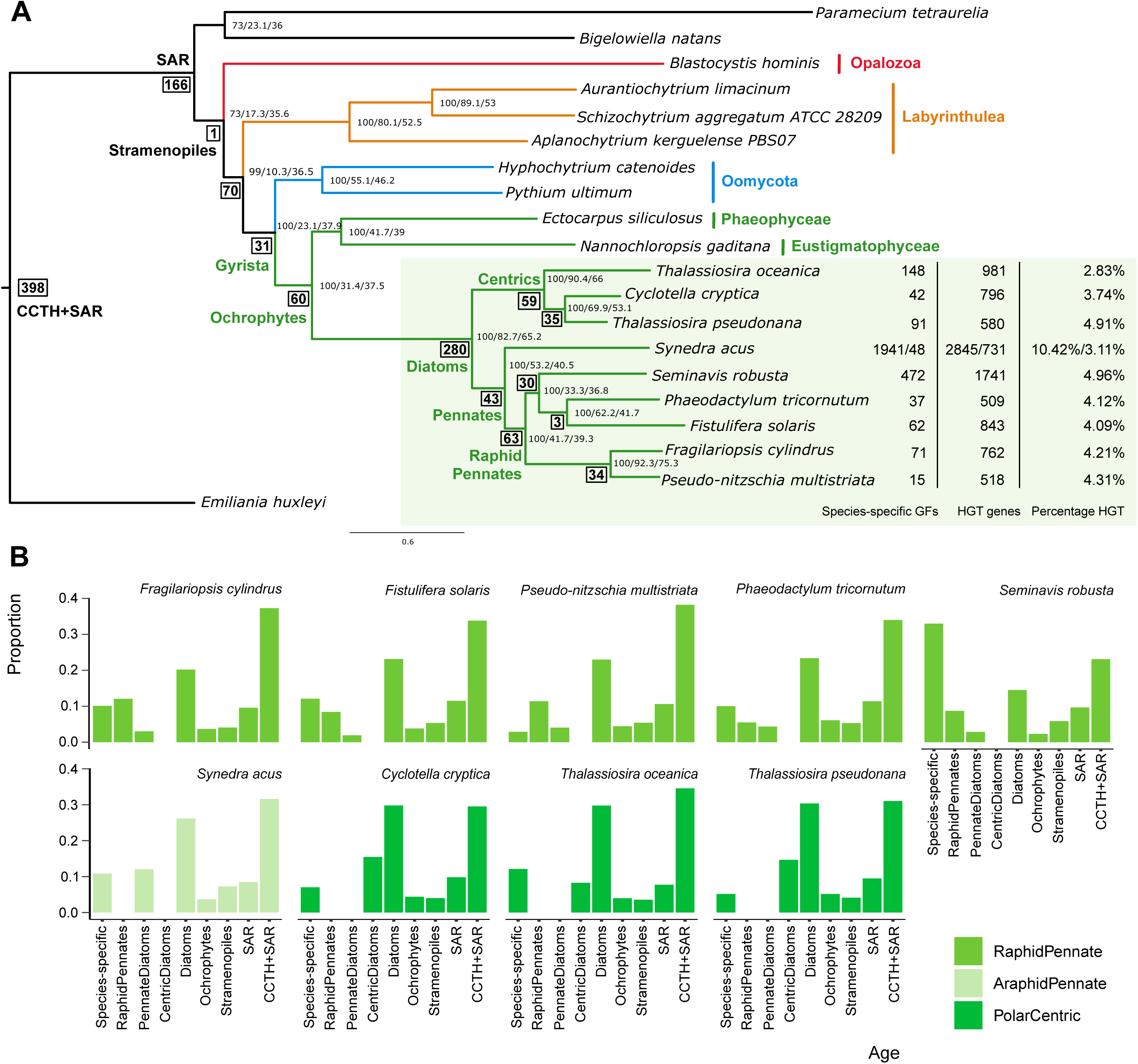
Overview of HGT events across diatoms. **a**, Species phylogeny determined by IQ-Tree with values at the internal nodes that denote bootstrap, gene- and site-concordance factors respectively. Branches are coloured according to their phylogenetic classification. Bold and framed values at internal nodes reflect the number of predicted HGT events. For diatoms, the number of species-specific GFs of HGT origin, the total number of HGT genes and its fraction of the proteome to be originating from HGT is tabularised. For *S. acus* the number of HGT genes both prior and after removal of contamination is mentioned. **b**, Distribution of the age classes of HGTs across the nine investigated diatom species.

To avoid the misclassification of contaminating DNA present in the genome assembly as genomic regions originating by HGT, several quality analyses were performed. The guidelines proposed by Richards and Monier^31^ were followed to exclude incorrect inference of HGT. Therefore, the gene origin was determined by phylogenetic tree construction followed by inspection of species-specific HGT genes. Also the percentage GC and the integration of HGT genes across chromosomes was assessed. First, the fraction of species-specific HGT was compared among all diatoms. More than 75% (2146/2844) of the predicted HGT genes in *S. acus* were only detected in this genome, while in all other species this fraction was drastically lower (11.58 +/- 9.25%) (Figure 1b). A donor analysis of these genes revealed that many were derived from *Sphingomonas sp*., which has been described to be associated with *S. acus* in culture^32^. Contigs flagged to be contaminant based on a nucleotide sequence similarity search against all available Sphingomonadales genomes were clearly separable from *S. acus* based on their significantly lower percentage GC (Figure 2a) (42.1% vs 63.3%, p-value <2×10^−16^). Therefore, all 695 nuclear contigs having a GC content above 50% were removed, reducing the nuclear *S. acus* genome size by 4 Mb to 94.38 Mb and retaining 23,719 genes. Interestingly, the HGT detection procedure succeeded in both flagging the contaminant as detecting HGT events in the *S. acus* genome (Figure 2b). Despite the fact that in several other diatoms the GC content was significantly different between genes from horizontal and vertical descent, the mean difference never exceeds two percentage points (Figure S1).

**Figure 2:**
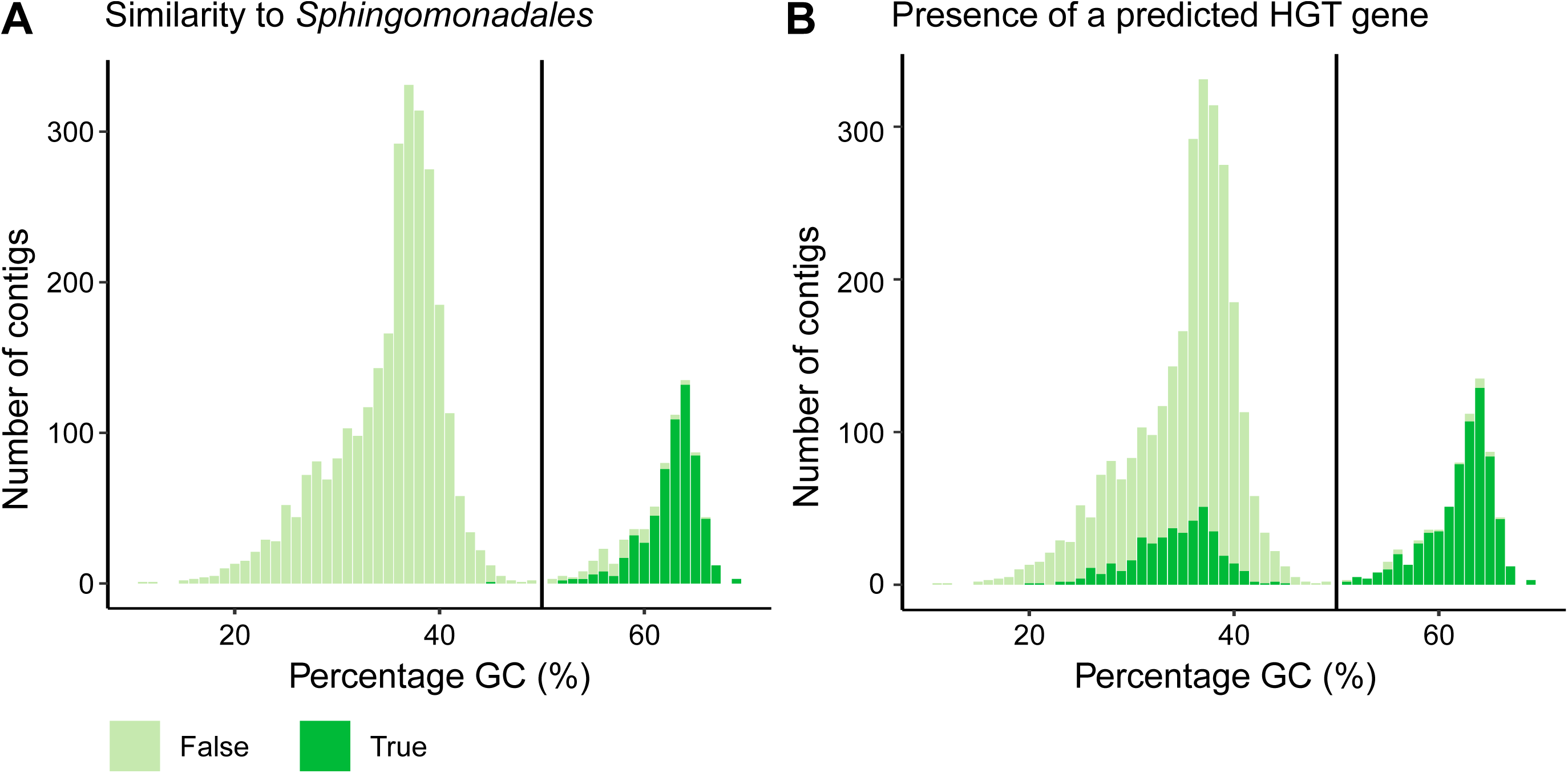
Contamination of Sphingomonadales in genome of *S. acus*. **a**, Percentage GC of *S. acus* contigs versus nucleotide sequence similarity to Sphingomonadales based on at least 70% identity and 25% alignment coverage. **b**, Percentage GC of *S. acus* contigs versus presence of a HGT candidate within each contig.

Next, the enrichment of HGTs per contig or chromosome was evaluated to assess whether certain regions are derived from contamination, yielding no clear examples of clustering of HGT genes on specific genomic locations. The distribution of HGTs across the chromosome-level genomes of P. tricornutum and *T. pseudonana* is plotted in Figure S2 and shows an unbiased distribution of HGT genes. As it has been proposed that the transfer of transposable elements could be associated with facilitating gene transfer^33^, the distance between every gene and its closest transposable element was calculated in *P. tricornutum*. Species-specific HGT genes were significantly closer to transposable elements (TEs) (p-value 1.6×10^−03^), while the same was also true for vertically descended species-specific genes (p-value 2.7×10^−14^). This suggests that novel genes are more likely to integrate and become fixed close to repetitive regions.

Except for a fraction of genes in the *S. acus* data set, we could not identify genomic properties indicating that the identified HGT genes are caused by contamination. In total, 7,461 diatom genes were defined as having HGT origin, covering 1,979 gene families. This reflects 509 to 1,741 genes per species, making 3 to 5 percent of the diatom gene repertoire predicted to be HGT (Figure 1a). This is similar to previous phylogenetic-based estimations of HGT content in diatoms, which ranged from 587 genes (4.8%) in *P. tricornutum*^7^ to 438 in *P. multistriata*^30^ (3.6%) and is slightly higher than what was reported in the anaerobic gut parasite *Blastocystis hominis*^34^ (2.5%), where next to bacterial HGT also other transfers were described. The lower frequency in this stramenopile could be due to its constrained and reduced genome size as a result of its parasitic lifestyle. On average, a HGT gene family consisted out of 3.76 diatom genes and 2.55 diatom species. In total, only 106 HGT families were present in all nine diatoms. For 69 gene families the HGT copies were significantly expanded in at least one species, of which notably 26 and 21 gene families were expanded respectively in *S. robusta* and *S. acus* (Table S2). Indeed, gene family expansion by duplication has been observed before following HGT integration in eukaryotes^1,35^ and this could be a strategy to diversify the original acquired function.

The age of all gene families of vertical descent was determined based on the lowest common ancestor of the observed species. Similarly, the most likely time point of integration for every HGT was determined using the species composition of the acceptor branch in the phylogenetic tree (Figure 1a). The large number of HGT gene families that can be attributed to the ancestor of diatoms is striking, ranging from 15% in *S. robusta* to 30% in *T. pseudonana* (Figure 1b). Another study^27^ also detected a continuous flux of genes from prokaryotes during the evolutionary history of *P. tricornutum*. However, they claimed that most influx occurred at ancestor of the photosynthetic Stramenopiles (Ochrophytes), while our results indicate this happened more recently in the ancestor of the diatom clade.

Finally, several structural gene features were evaluated according to their mode of inheritance. The coding gene length of vertically descended species-specific genes in all diatom species was significantly shorter compared to all other genes (p-value <2×10^−16^) and significantly shorter to the species-specific HGT genes in all diatoms, except for *T. pseudonana* (Figure S3). In yeast, it has also been observed that *de novo* genes were on average shorter than conserved and horizontally transferred genes^36^. Species-specific HGT genes on the other hand, were significantly shorter to all other genes in *C. cryptica* (p-value 2.1×10^−02^) and *S. robusta* (p-value 1.4×10^−05^). Given that introns are a typical eukaryotic gene feature, HGT genes are expected to have a shorter total intron length, especially for recent acquisitions as HGT genes adapt to their recipient genome. The intron length of HGT genes was significantly shorter in several pennate diatoms (*F. solaris*: 1.8x×10^−03^, *P. tricornutum*: 3.4×10^−02^, *S. robusta*: 1.1×10^−07^ and *S. acus*: 4.6×10^−03^) and for several diatoms the young species-specific HGT genes had shorter introns than the rest of the gene repertoire (*F. cylindrus*: 1.1×10^−03^, *F. solaris*: 9.8×10^−03^, *P. tricornutum*: 1.1×10^−02^, *S. robusta*: 2×10^−09^ and *C. cryptica*: 4.9×10^−02^). These results indicate that introns become an emerging property of HGT genes after integration.

### The functional landscape of diatom HGT genes

To gain insight in the functional repertoire of HGT genes, a gene ontology (GO) and functional domain (Interpro) enrichment was performed. Out of the 7,461 diatom HGT genes, 6,024 (81%) were annotated with an Interpro domain and 3,893 (52%) with a GO term. The only GO term which was enriched for HGT genes in all nine diatom species is pseudouridine synthesis (GO:0001522), while enriched protein domains covered pseudouridine synthase (IPR006145), S-adenosyl-L-methionine-dependent methyltransferase (IPR029063) and nitroreductase (IPR029479). An overview of the enriched functional categories across different ages can be found in (Figure S4, Figure S5). A more in-depth exploration of several functional categories is given in Supplementary Note 1, while an overview of all discussed functions and their corresponding gene families can be found in Table S3.

### Cobalamin uptake

Cobalamin (vitamin B12) is a complex molecule composed out of a central cobalt-containing corrin ring, a lower ligand of 5,6-dimethylbenzimidizole (DMB) and an upper axial ligand that can either be an hydroxy-, cyano-, methyl or adenosyl group. Vitamin B12 acts as a coenzyme in three enzymes in eukaryotes: methylmalonyl-CoA mutase (MCM), type II ribonucleotide reductase (RNRII) and methionine synthase (METH). Despite that more than half of the algal species surveyed (171/326)^37^ are auxotrophic for vitamin B12, including 37 out of 58 diatoms, *de novo* synthesis has only been described to occur in prokaryotes. Therefore cobalamin availability alters the composition of marine phytoplankton communities^38^. The exchange of cobalamin in return for organic compounds is believed to underpin the close mutualistic interactions between heterotrophic bacteria and auxotrophic algae^39^ A correlation was detected between the scattered phylogenetic pattern of absence of a cobalamin-independent methionine synthase (METE) and auxotrophy for this vitamin^40^. It has been suggested that this loss has a biogeographical basis as there is a tendency for diatoms occurring in the Southern Ocean to retain METE more often^41^. Moreover, it has been recently proven that cyanobacteria produce the chemical variant pseudo-cobalamin, where adenine substitutes DMB as the lower ligand, which is less bioavailable to eukaryotic algae^42^. However, some species, including *P. tricornutum* and *E. huxleyi*^43^ can remodel this to cobalamin using CobT, CobS and CobC via the nucleotide loop assembly^39,42^. Here BluB, necessary for DMB production^44^, was detected to have originated by HGT from alphaproteobacteria in *F. cylindrus, P. multistriata* and *P. multiseries* (Figure 3a). More than 90 percent of the cobalamin-producing alpha- and gammaproteobacteria encode BluB^39^. Moreover, five HGT genes were detected in the final synthesis of the cobalamin biosynthesis pathway, which can also function as scavenging and repair genes: CobN, CobA/CobO, CobQ/CbiP, CobD/CbiB and CobU/CobP (Figure 3a). These genes were previously also detected in diatom metatranscriptomes and *P. granii*^45,46^, where CobN, CobS and CobU were more highly expressed under iron replete conditions. Interestingly, for all diatoms, except for *C. cryptica*, the CbiB gene also contains the CbiZ domain, which is involved in the removal of the lower ligand^43^. Only *F. cylindrus* and *P. multiseries* contain the full suite of these detected HGT genes in their cobalamin pathway, while *P. tricornutum, F. solaris* and *S. acus* possess none (Figure 3a). Interestingly, while *P. multiseries* is auxotrophic for cobalamin, *F. cylindrus* is not. Thus, despite the presence of the cobalamin independent methionine synthase METE, *F. cylindrus* expanded its repertoire of cobalamin synthesis genes and prefers to maximally optimize its uptake to perform methionine synthesis by the more efficient METH. By querying the metatranscriptomic TARA Oceans data it was clear that CobU, CbiZ+CbiB and CobQ are significantly correlated with nitrate concentration and day length (Figure 3b) (Figure S6,Figure S7), while CobU and CbiZ+CbiB are anti-correlated with temperature (Figure S8) and CobU and CobQ are anti-correlated with iron (Figure S9). HGT genes in the cobalamin pathway are particularly abundant in the Southern and Pacific Ocean (Figure 3b). The lower production rate of bacteria in low temperature and the photodegradation of cobalamins, which could be of particular importance during arctic summers, might explain the cobalamin limitation and the specific expression of vitamin B12-related genes in these regions of the ocean.

**Figure 3:**
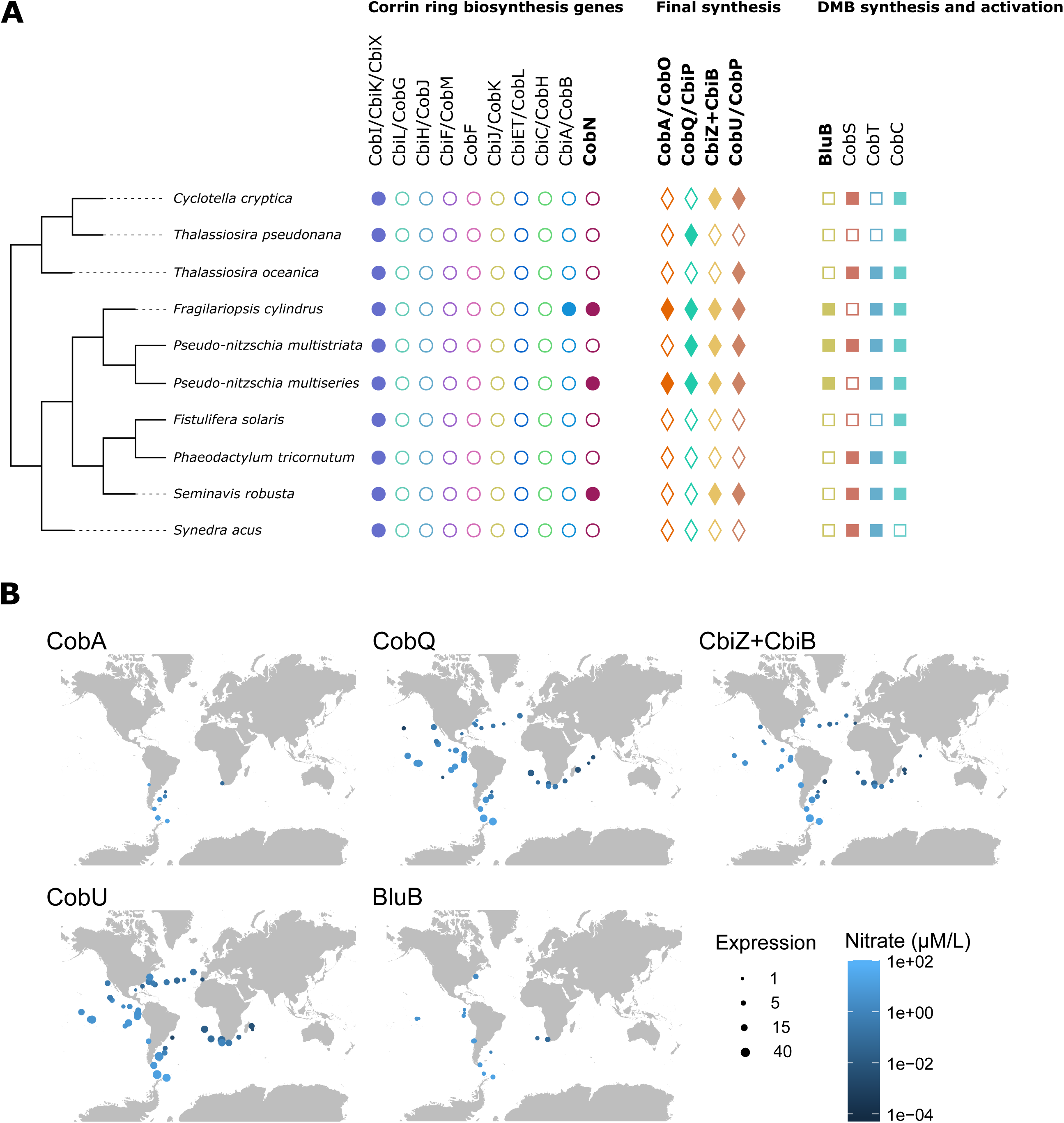
Cobalamin pathway in diatoms. **a**, Overview of the cobalamin biosynthesis pathway and its presence in diatoms. Genes in bold are of horizontal descent and the presence of a gene is displayed by a filled circle, diamond or square depending on its position in the pathway. **b**, Expressed number of diatom sequences of several HGT genes involved in the cobalamin pathway across stations sampled worldwide during the TARA Oceans project in the surface layer, coloured according to their nitrate concentration.

### Environmental adaptation to light sensing and cold protection

Diatoms employ photosensory proteins to gain information about their environment and respond to changing light conditions. Proteorhodopsins (PR) perceive light to drive ATP generation and are especially important when photosynthesis is comprised during iron-limiting conditions. This study confirms the bacterial origin of the PR-genes in *F. cylindrus* and *Pseudo-nitzschia granii*^47^, next to brown algae, dinophytes and haptophytes. Furthermore, the red/far-red light sensing phytochrome DPH1^48^ was detected as HGT in *P. tricornutum* (1 copy), *S. robusta* (4), *S. acus* (7), *C. cryptica* (1) and *T. pseudonana* (1), and formed together with brown algae an independent branch from green algal and fungal DPHs, similar as in previous reports^48,49^ and was predicted to have originated from HGT.

Arctic diatoms such as *F. cylindrus* undergo periods of prolonged darkness, low temperature and high salinity. Their ability to thrive in these conditions could be partially attributed to cryoprotectants that interfere with the growth of ice^50^. Ice-binding proteins were found to be laterally transferred from a basidiomycete lineage to *Fragilariopsis curta* and *F. cylindrus*^51^. Also the phylogenetic tree inferred in this study detected relatedness between fungal and diatom antifreeze proteins, but wasn’t classified as HGT as this pipeline only detects bacterial-to-eukaryotic HGT. However, a second gene family of *F. cylindrus* proteins containing the ice-binding protein domain (IPR021884), was found to be transferred from *Cryobacterium*.

### Carbon and nitrogen metabolism

Diatoms can rapidly recover from prolonged nitrogen limitation due to presence of the urea cycle that allows for carbon fixation into nitrogenous compounds^52^. Two genes in the metabolic branches derived from this pathway, carbamate kinase and ornithine cyclodeaminase were found to be laterally transferred, both here as in previous studies^7,52^. The latter enzyme is responsible for the conversion of ornithine to proline, which is the main osmolyte during salt stress in diatoms. Another way of a nitrogen storage and translocation is the catabolism of purines to urate that can be further degraded to allantoin. It was found that plants and diatoms independently evolved a fusion protein (Urah-Urad domain; allantoin synthase) to perform the second and third step in this urate degradation pathway^53^. Exactly as in^53^, this gene was detected to be laterally transferred from alphaproteobacteria, where this fusion event occurred, to the ancestor of haptophytes and stramenopiles.

Moreover, several genes in carbohydrate metabolism were found to be laterally transferred. The acetyl-CoA conversion to acetate occurs in a two-step process where phosphate acetyltransferase (PTA) adds a phosphate group to form acetylphosphate, that is in turn is catalyzed to acetate by acetate kinase (ACK)^54^. The PTA gene family was found to have bacterial origins and emerged in the ancestor of haptophytes and stramenopiles. In all diatoms, except for *F. cylindrus* and *P. multistriata*, multiple copies were found of this gene. Also acetate kinase was detected as a HGT gene in the pennate diatoms *P. tricornutum, S. robusta* and *S. acus*. Furthermore, this enzyme was predicted to be involved in the bifid shunt^54^. Here, the key enzyme XPK cleaves xylulose-5-phosphate to acetyl-phosphate and glyceraldehyde-3-phosphate, followed by conversion of acetyl-phosphate to acetate by ACK. Also XPK was laterally transferred in the pennate diatoms, single-copy in *P. tricornutum* and significantly expanded to five copies in *S. acus*. XPK and ACK are syntenic in *P. tricornutum*, what was already suggested to point to a bacterial origin as this spatial organization is also detected in Proteobacteria and Cyanobacteria^54^. Interestingly, *S. acus* has also conserved the physical association of XPK and ACK and maintained a bidirectional promoter, although an inversion of the gene order occurred (Figure S10). Furthermore, phosphofructokinase and the cytosolic fructose-bisphosphate aldolase Fba4 in the glycolysis^55^, phosphopentose epimerase^56^ in the pentose phosphate pathway and a putative D-lactate dehydrogenase are enzymes that were predicted to be transferred from bacteria present in diatoms. Finally, also bacterial xylanases, glucanases and glucosidases expanded the carbohydrate metabolic repertoire in diatoms.

The biosynthetic aspartate-derived pathway to synthesize the four amino acids, lysine, threonine, methionine and isoleucine was completed due to HGT^57^. Aspartate semialdehyde dehydrogenase (asd) performs the second step in this pathway and is derived from Proteobacteria. The end product L-aspartate 4-semialdehyde can either be used by dihydrodipicolinate synthase (dapA) towards lysine biosynthesis, or by homoserine dehydrogenase (thrA) towards threonine and methionine. Both genes were laterally transferred from bacteria. The metabolic pathways of other amino acids were also affected, the last step in tryptophan synthesis is achieved by tryptophan synthase. While in diatoms the alpha and beta subunit of this enzyme are merged, in *P. tricornutum* an extra copy of the beta subunit is present^58^ (Phatr3_J52286) that was deemed bacterial. Also alanine racemase, arginine biosynthesis ArgJ, leucyl-tRNA synthetase leuRS2, glycyl-tRNA synthetase glyRS2 and tyrosine-tRNA ligase tyrRS2 were laterally transferred.

### Selection pattern for HGT genes in *P. tricornutum*

Genomic sequence information from ten *Phaeodactylum* accessions, belonging to four clades sampled across the world^59^, was used to do determine the maintenance and selection pattern across the detected HGT genes. The retention of species-specific HGT genes across different strains confirmed their horizontally derived origin and did not point to contamination (for more details, see Supplementary Note 2). Moreover, analyzing gene selection patterns gives an indication on the strength of functional conservation. Variant calling resulted in a data set of 585,715 high-confidence bi-allelic SNPs. The total number of SNPs per strain across the genome was low and ranged from 0.96% to 1.37% (Table S4). To detect selective pressure, πN/πS, was calculated. This metric compares the fraction of synonymous and nonsynonymous mutations within a coding open-reading frame across strains. A gene experiencing neutral evolution has a πN/πS value of 1, whilst a value smaller than 1 signifies negative purifying selection. The smaller the ratio of non-synonymous and synonymous nucleotide diversity, the stronger is the level of purifying selection acting on the gene. The average synonymous nucleotide diversity (πS) across all accessions is 0.009, while the non-synonymous nucleotide diversity (πN) is 0.003, thus the genome-wide average πN/πS ratio is 0.3. This value is similar to what was described in Rastogi et al.^59^ and means most genes undergo strong purifying selection. The average πN/πS for genes of vertical descent is 0.302, while for HGT genes it is significantly lower at 0.268 (p-value 5.9×10^−4^). When comparing the πN/πS ratios for HGT and vertical genes across age classes (Figure 4), it is apparent that this difference is due to the youngest gene categories, being the *Phaeodactylum*-, raphid pennate diatoms and pennate diatoms specific genes, where vertical genes are less constrained than HGT genes in those age categories. To the best of our knowledge, this is the first time the selection pressure of bacterial HGT genes is assessed in unicellular eukaryotes and compared with vertically descended genes while taking age into account. Although it has already been observed that *de novo* genes display patterns of rapid evolution and the strength of purifying selection increases with age^36^, it is remarkable to observe that HGT genes deviate from this pattern. Unlike recent innovations from vertical descent, young HGT genes are quickly integrated in the biological network exemplified by their high levels of purifying selection.

**Figure 4:**
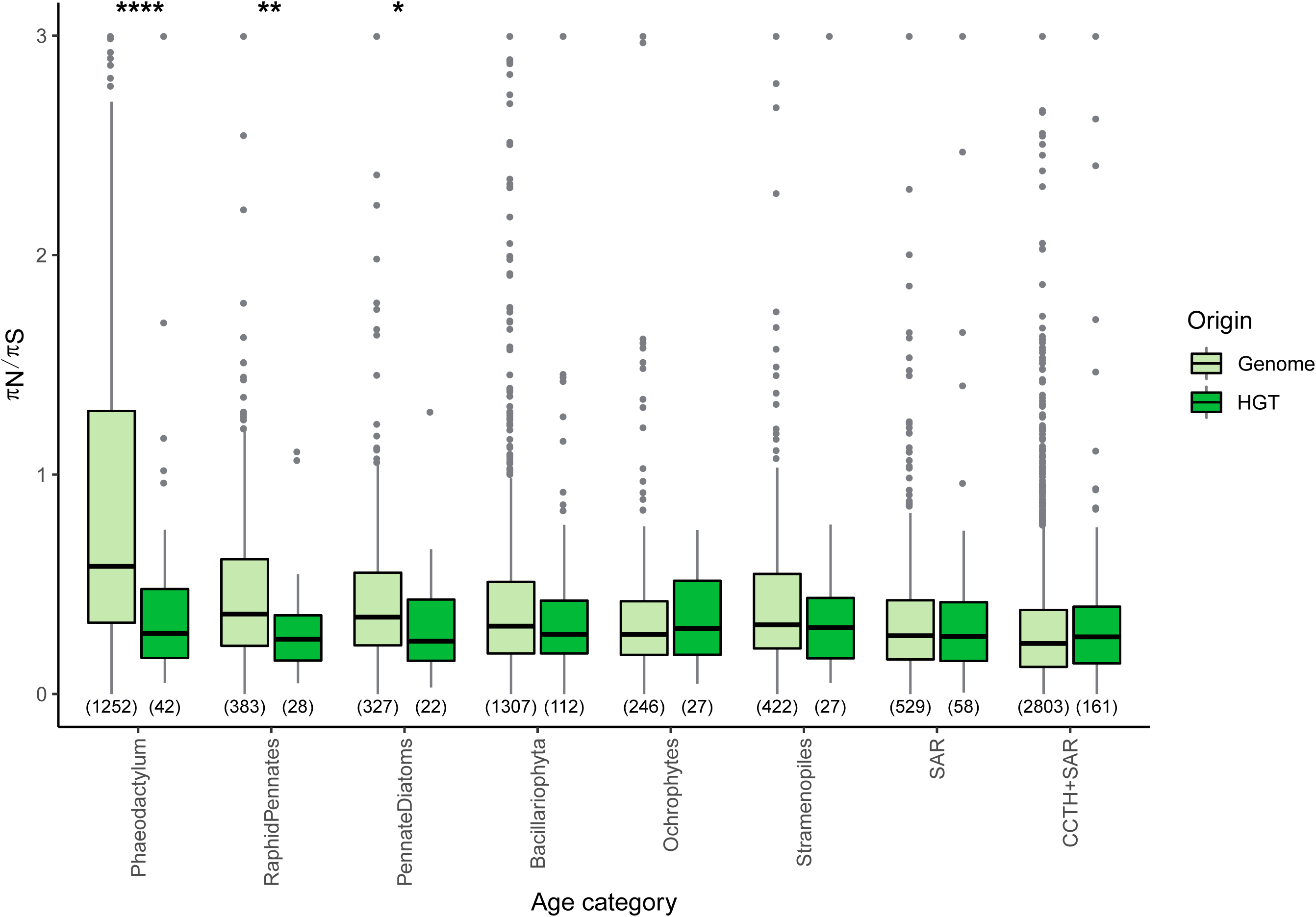
Selective pressure over time in *P. tricornutum*. Distribution of selective pressure, measured by πN/πS, across age classes sorted from young to old and per origin in *P. tricornutum*. Number of genes is indicated in between brackets. The asterisks denote a statistical difference per type within the same age category and have the following confidence range for p-values; * : 0.05, ** : 0.01, *** : 0.001, ****: 0.0001.

### Expression and co-expression network analysis of HGT genes

The availability of RNA sequencing experiments in several diatoms, allows for the construction of genome-wide gene expression atlases quantifying gene expression levels across a wide range of conditions. These compendia consisted out of 13 to 76 conditions per species, all having biological replicates per condition (Table S5). The vast majority of HGT genes are expressed: in *P. tricornutum* all HGT genes are expressed, in *T. pseudonana* 558 out of 580 (96%), in *S. robusta* 1597 out of 1741 (92%) and in *F. cylindrus* 741 out of 762 (97%). Given that most HGT genes are kept under purifying selection in *P. tricornutum* and are transcribed in diatoms, this is indicative that they are functional and can play a vital role in expanding the functional repertoire. Indeed, 64% of the predicted HGT genes in *P. tricornutum* were translated into proteins in a proteogenomic analysis^60^. This is similar to 63% of all proteins in the genome that were detected to be translated.

Next, the expression specificity was calculated per gene, where a low value signifies broad expression in many (or even all) conditions and a value close to one indicates expression in one or a few. Species-specific genes have a higher mean condition-specific expression, both for vertical as horizontal derived genes in *P. tricornutum* (p-value <2×10^−16^, 1.1×10^−06^), *S. robusta* (<2×10^−16^, 1.6×10^−13^), *F. cylindrus* (<2×10^−16^, 2.2×10^−03^) and *T. pseudonana* (<2×10^−16^, 3.7×10^−02^). A declining trend of condition-specificity was observed over time. Whenever there was a significant difference in condition specificity between HGT and vertical genes within the same age category, HGT genes consistently displayed on average a more specific expression pattern (Figure 5). A high tissue-specificity for species-specific genes which decreases over time has also been observed in mouse^61,62^. Interestingly, the selection pressure in *P. tricornutum* across all age classes and per mode of inheritance is not correlated with expression specificity (Figure S11), showing that genes having a highly specific expression are not necessarily under less purifying selection, defying the trend that was previously observed in mammals^63^.

**Figure 5:**
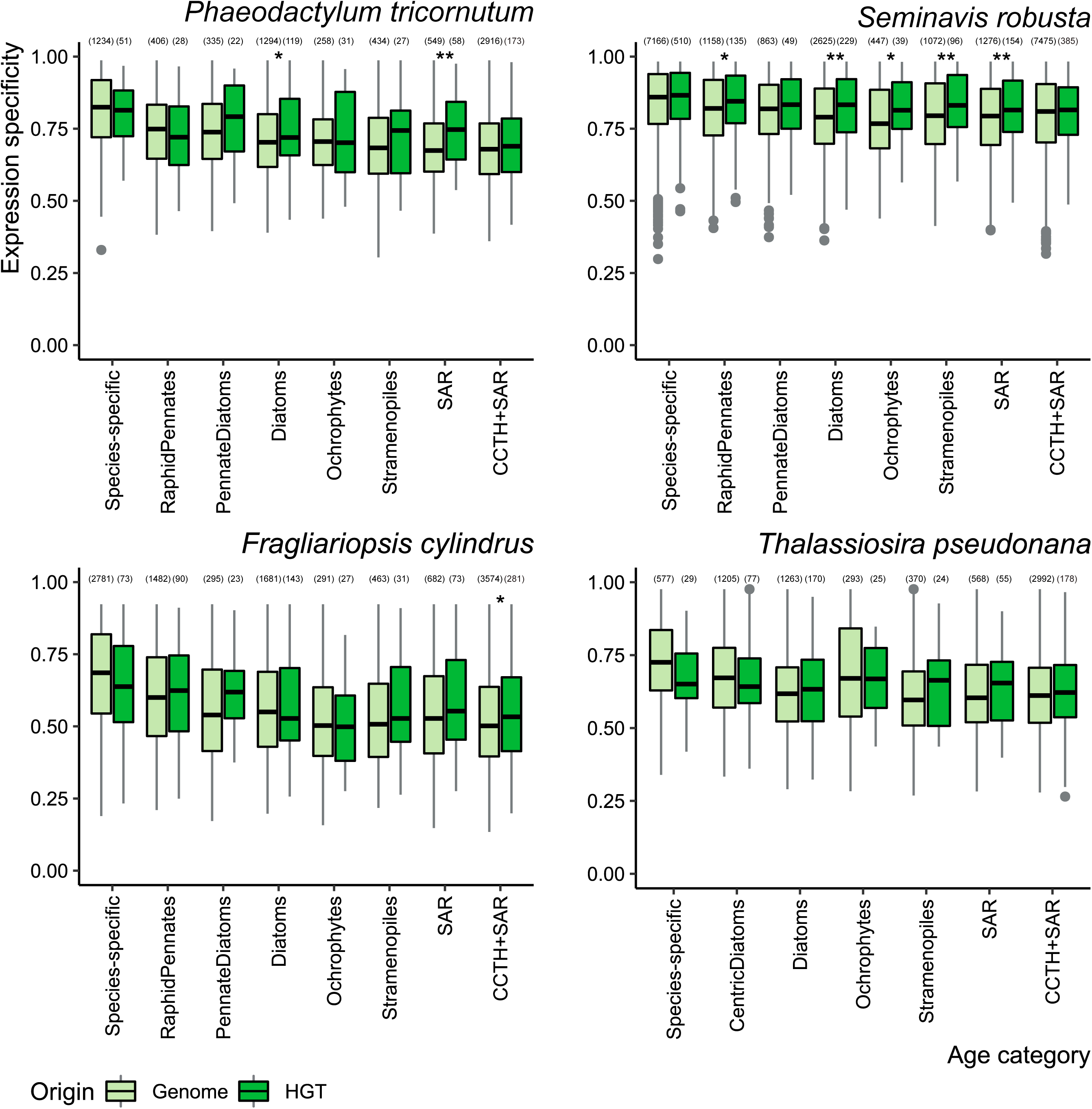
Expression specificity over time in diatoms. Distribution of expression specificity across age classes sorted from young to old and per origin in four diatoms. The number of genes is indicated in between brackets. The asterisks denote a statistical difference per type within the same age category and have the following confidence range for p-values; * : 0.05, ** : 0.01, *** : 0.001, ****: 0.0001.

Based on a global co-expression *P. tricornutum* network constructed using an expression atlas comprising 211 samples, for every gene in *P. tricornutum* the co-expression neighbourhood was defined as a module. Subsequently, these modules and the known gene function for genes part of this module were used, through guilt-by-association analysis, to gain functional insights in the detected HGT genes. For 19 HGT genes the co-expression modules confirmed enrichment for at least one known function. Fructose-bisphosphate aldolase Fba4 is enriched in its co-expression module for genes involved in a carbohydrate metabolic process and aspartate semialdehyde dehydrogenase (asd) has enrichment for amino acid biosynthesis. New functions were attributed to 320 out of 509 HGT genes based on significant GO enrichment of the co-expression modules. For example, two HGT proteins involved in amino acid synthesis - tryptophan synthase β chain (p-value 2.2×10^−16^) and ArgJ (p-value 1×10^−15^) - are predicted to be coregulated with photosynthetic genes, although both proteins do not contain the chloroplast targeting peptide. The co-expression neighbourhood of phosphofructokinase is significantly enriched (p-value 3.7×10^−06^) to be involved in the Krebs cycle, while this protein is part of the glycolysis and directly upstream of the citric acid cycle. Also Phatr3_J40382, which contains a pyruvate kinase-like domain, is enriched for this GO term, and this corroborates its metabolic function. Methionine sulfoxide reductase MsrB (Phatr3_J13757) was predicted to have a similar expression pattern to genes partaking in iron-sulphur cluster assembly (p-value 1.1×10^−03^) and metal ion transport (p-value 1.4×10^−02^). In yeast these proteins were already shown to have a protective role for FeS clusters during oxidative stress^64^. The far-red light phytochrome DPH1 is enriched for genes involved in transcriptional regulation (p-value 5.5×10^−03^), which could point to its primary role in the light sensing cascade. These results demonstrate that co-expression network analysis offers a pragmatic means to predict the biological processes HGT genes are involved in.

## 4. Discussion

Through the application of phylogeny-based HGT detection, we identified 1,979 gene families with a horizontal origin in diatoms. Although HGT detection has been previously performed in diatoms, this is the first large-scale and systematic approach of HGT detection across all available sequenced diatoms. While some previous studies were based on phylogenetics^7,30^, most relied purely on sequence homology searches^24,25,27^, while it has been shown that the degree of gene similarity does not necessary necessarily reflect phylogenetic relationships^65,66^. Although HGT had been previously predicted in *P. multistriata, C. cryptica, T. oceanica* and twice in *P. tricornutum*, only a fraction of HGT genes were confirmed across these studies, going from 6 to 45% (Figure S12). This could be due to the usage of different methods, criteria and underlying databases. For example, horizontal genes were defined in *T. oceanica* if they didn’t show high similarity with any other stramenopile and thus contain *T. oceanica*-specific genes from both bacterial- and eukaryotic-to eukaryotic origin. Nonetheless, the overlap between all methods is still significantly higher than expected by chance for all species.

Assessing the strength of purifying selection using genome-wide nucleotide diversity information showed that lateral gene transfer is quickly followed by fixation compared to the retention of new genes originating from non-coding regions, so-called *de novo* genes. Moreover, most species-specific HGT genes are present across all strains in *P. tricornutum*, suggesting they were acquired prior the divergence of these strains and are actively maintained in the population. This underlines the importance of HGT in diatoms and indicates that fixation of a laterally transferred gene takes place quickly after the initial uptake. Indeed, in grasses it was recently shown that several plant-to-plant LGT fragments were rapidly integrated and spread across the population, after which erosion occurred on neutrally selected genes within those fragments^67^.

Among the HGT events, we detected the transfer of five concurrent genes of the vitamin B12 biosynthetic pathway. A cobalamin addition experiment in a high-nutrient low chlorophyll (HNLC) region in the Gulf of Alaska significantly altered the species composition, going from diatom-dominated plankton to an increased fraction of ciliates and dinoflagellates^68^. This could be explained by the presence of these HGT genes and the corresponding enhanced uptake mechanism of vitamin B12 and its analogues, which give diatoms a competitive advantage during limiting conditions. In conclusion, our results support a high genetic plasticity and ability for local adaptation in diatoms due to HGT.

## 6. Material and methods

### Gene family construction

The publicly available genomes of 17 stramenopiles, 1 alveolate, 1 rhizarian and 1 haptophyte (listed in Table S1) were downloaded and their nuclear proteomes, totalling to 398,001 protein-coding genes, were searched for similarity in an all-against-all fashion with BLASTp (version 2.6+) using an e-value cut-off of 10^−5^ and retaining maximum 4,000 hits. Next, clustering of these protein-coding genes was performed using OrthoFinder (version 2.1.2)^69^.

### Species tree phylogeny

To delineate the species phylogeny for all SAR members, using *Emiliania huxleyi* as an outgroup, OrthoFinder gene families where all species have a copy number of either 1 or 2 genes were selected, and one gene sequence was randomly picked in case of duplication. MUSCLE^70^ was used to build a concatenated sequence alignment. Afterwards IQ-Tree (version 1.7.0b7)^71^ was used to build a concatenated tree using 1,000 bootstraps, estimate the single-locus trees and finally to calculate the gene- and site-concordance factors of the inferred species tree^72^.

### HGT detection

The NCBI non-redundant protein database (download date 08/06/2018) was complemented with the proteomes of 20 species (Table S1). Diamond (version 0.9.18.119)^73^ searches were performed in sensitive mode against this database for all proteins of these 20 species, retaining maximum 1,000 hits per query. Hits were reduced to maximum five sequences for each order and 15 sequences per phylum. Genes families with at least one copy in a diatom and at least one third of the diatom members having a bacterial hit were analysed. The hits of all diatom members were combined and clustered using CD-HIT (version 4.6.1)^74^ based on a 95% identity cut-off. Next, the sequences were aligned with MAFFT (version 7.187)^75^ in automatic mode. Maximum likelihood trees are produced using IQTree (version 1.6.5)^71^ including a test for the best fitting protein model (-mset JTT, LG, WAG, Blosum62, VT, Dayhoff)^76^. The FreeRate model was used to account for rate heterogeneity across sites (-mrate R), empirical base frequencies were calculated (-mfreq F) and 1,000 rounds of ultra-parametric bootstrapping (-bb 1000) (UFBoot2)^77^ were run.

Phylogenetic trees were reordered based on midpoint rooting, unless the whole eukaryotic fraction formed a cluster and then this cluster was used as a subtree for rooting. For every node having a bootstrap support ≥ 90, and consisting out of a bacterial and eukaryotic subtree the last common ancestor (LCA) was defined. When the eukaryotic subtree consisted out of more than 20 sequences, they could only compose at most 85% of the total number of sequences belonging to that node. When several nodes complied to these rules having the same eukaryotic fraction, the last common ancestor of the bacterial subset was considered the donor of this event. To avoid classifying endosymbiotic gene transfer (EGT) incorrectly as HGT, only bacterial-to-eukaryotic events were analysed and events older than SAR + Haptophytes, also dubbed SAR+CCTH, were discarded.

### Gene family expansion

Expanded families were delineated by calculating the Z-score profile of the gene copy number per HGT family across all diatoms excluding the allodiploid *F. solaris*. Families where the variance is larger than two and the Z-score for a particular species is larger than three, were deemed expanded in that species.

### Contamination detection

Structural genomic annotation features for sequenced diatoms were retrieved and GC content, coding sequence length, number of introns per gene and intron length were compared between horizontally and vertically transferred genes among several age categories. Also the distance for every gene in *P. tricornutum* to the closest transposable element as defined by ^27^, centromeric and telomeric regions elucidated by ^78^ was calculated and compared among the different origins. Statistical significance was calculated by the Wilcoxon rank sum test. For the diatoms *T. pseudonana* and *P. tricornutum*, whose genomes are resolved on chromosome-scale level, the distribution of HGT genes was plotted using R.

To assess the degree of contamination from *Sphingomonas sp*. in *S. acus*, 914 genomes of the order *Sphingomonadales* were retrieved from NCBI and a nucleotide blast against the *S. acus* genome was performed. Contigs having at least 70% identity and 25% alignment coverage were deemed to have *Sphingomonadales* origin.

### Functional interpretation of HGT genes

The proteomes of all species were functionally annotated using Interproscan (version 60)^79^ in order to obtain functional domain annotations and Gene Ontology (GO) terms. KEGG orthology identifiers^80^ were attained using EggNOG-mapper^81^. For all diatoms, the chloroplast targeting signal was predicted using ASAFind (version 1.1.7)^82^. Only GO terms within the subtree ‘biological process’ were taken into account. These terms were expanded to also contain all ancestral functional information. GO and Interpro domain enrichment was performed on the HGT genes per species using hypergeometric testing, and multiple hypothesis testing was constrained using Benjamini–Hochberg correction (q value < 0.05). Functional enrichments found in at least two species were visualized using the ComplexHeatmap package^83^ (R version 3.4) and clustered using the complete linkage method.

Tandem duplicates were defined as genes belonging to the same gene family and located within 15 genes of each other and identified using i-ADHoRe v3.0^84^ (alignment method: gg2, gap size 15, tandem gap 15, cluster gap 15, q-value 0.85, probability cut-off 0.01, anchor_points 3, level_2_only FALSE, FDR as method for multiple hypothesis correction).

### Metatranscriptome analysis

For several selected HGT gene families involved in cobalamin synthesis a HMM profile was created using hmmer3 v3.1b2^85^ and these were uploaded to the Ocean Gene Atlas webserver^86^ to query the eukaryotic MATOU gene dataset^87^ (blastp, evalue cut-off 10^−10^) linked the metatranscriptomic TARA Oceans data. Only sequences taxonomically assigned as diatoms were further analysed. The abundance was estimated as the number of sequence per station and depth.

### Population genetics

Data from ten resequencing strains was downloaded from the public repository SRA (https://www.ncbi.nlm.nih.gov/sra) (SRR6476693-SRR6476702) and these reads were mapped to the *Phaeodactylum tricornutum* genome using BWA-mem (version 0.7.17)^88^. The read alignments per strain were filtered to only include unique mappings without chimeric alignments using samtools (version 1.6)^89^. SGSGeneLoss (version 0.1)^90^ was run in a relaxed mode to determine to presence/absence pattern of all genes across the ten strains (minCov=1, lostCutoff=0.05, thus requiring only one read and 5% gene coverage to be perceived as present). The resulting phylogenetic pattern of HGT genes was visualized using the ComplexHeatmap package^83^ (R version 3.4) and clustered using the complete linkage method.

SNP calling was performed per strain using HaplotypeCaller, after which SNPs were integrated using GenotypeGVCFs. Both methods are available within the GATK framework (version 3.7)^91^. SNPs were filtering using the GATK recommended hard filters (QD<2.0; FS>60.0; MQ<40.0; MQRankSum ≤ 12.5; ReadPosRankSum ≤ 8.0)^92^ and only bi-allelic SNPs were retained.

To estimate the degree of negative purifying pressure across the proteome, only coding positions having a read depth ≥ 10 across all strains were considered, calculated using SAMtools mpileup^93^. In total, 89% of all genic positions could be analysed and 272,235 SNPs were observed in these regions. We used SnpEff (version 4.3t)^94^ to predict the individual effect per SNP and πN/πS was calculated taking only the callable positions for complete codons into account and correcting for the allele frequency of the mutation in the population. Statistical significance of difference in selective pressure across mode of inheritance and age classes was calculated by the Wilcoxon rank sum test.

### Expression and co-expression analysis

An expression atlas for every diatom species which has RNA-Seq expression data available was generated. First, relevant experiments were searched using Curse^95^, which also allows the user to identify and curate replicates. The experiments listed in (Table S5) were used to generate the expression compendia. Next, the atlas was generated using Prose^95^, which uses the SRA toolkit to download the raw data locally, FastQC to perform quality control and adapter detection, Trimmomatic for automatic read trimming and finally kallisto for expression quantification in transcripts per million (TPM). Genes were deemed expressed when having a TPM value of at least 3. The condition-specificity, also known as tau^96^, of every gene was calculated as follows, where x is the TPM value per condition, max is the maximal expression of a gene and n is the number of conditions in the expression compendium:

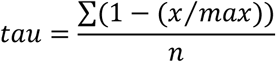

Condition-specific genes were defined as having a tau value of bigger or equal than 0.9.

The generated expression atlas for *P. tricornutum* was also used to define co-expression clusters. The Pearson correlation was calculated in a pairwise manner between all genes and the highest reciprocal rank (HRR)^97,98^ was determined at 23, by maximizing the recovery of known GO annotations, while restraining the number of novel predictions. A cluster was defined for every gene based on this cut-off and GO enrichment using hypergeometric testing was run per cluster. Multiple hypothesis testing was constrained using Benjamini–Hochberg correction (q value < 0.05).

### Data availability

All gene families, phylogenetic trees of horizontal descent and the dating of the HGT events within these gene families are available on Zenodo (https://zenodo.org/record/3555201).

## 7. Author contributions

E.V. wrote the manuscript, performed species topology delineation, HGT detection analysis, functional interpretation, metagenomic and population genomic analysis. T.D. aided in HGT delineation and performed co-expression analysis. C.M.O-C. aided in population genomic analysis and performed expression analysis generation of *S. robusta*. K.V. supervised the project. All authors read, edited and approved the manuscript.

## 8. Acknowledgments

E.V. wants to acknowledge the funding obtained by the BOF project GOA01G01715.

