## Supplementary material for "Systematic and functional analysis of horizontal gene transfer events in diatoms": SupplemtentalMaterial

### Supplemental material

#### Supplementary Figures

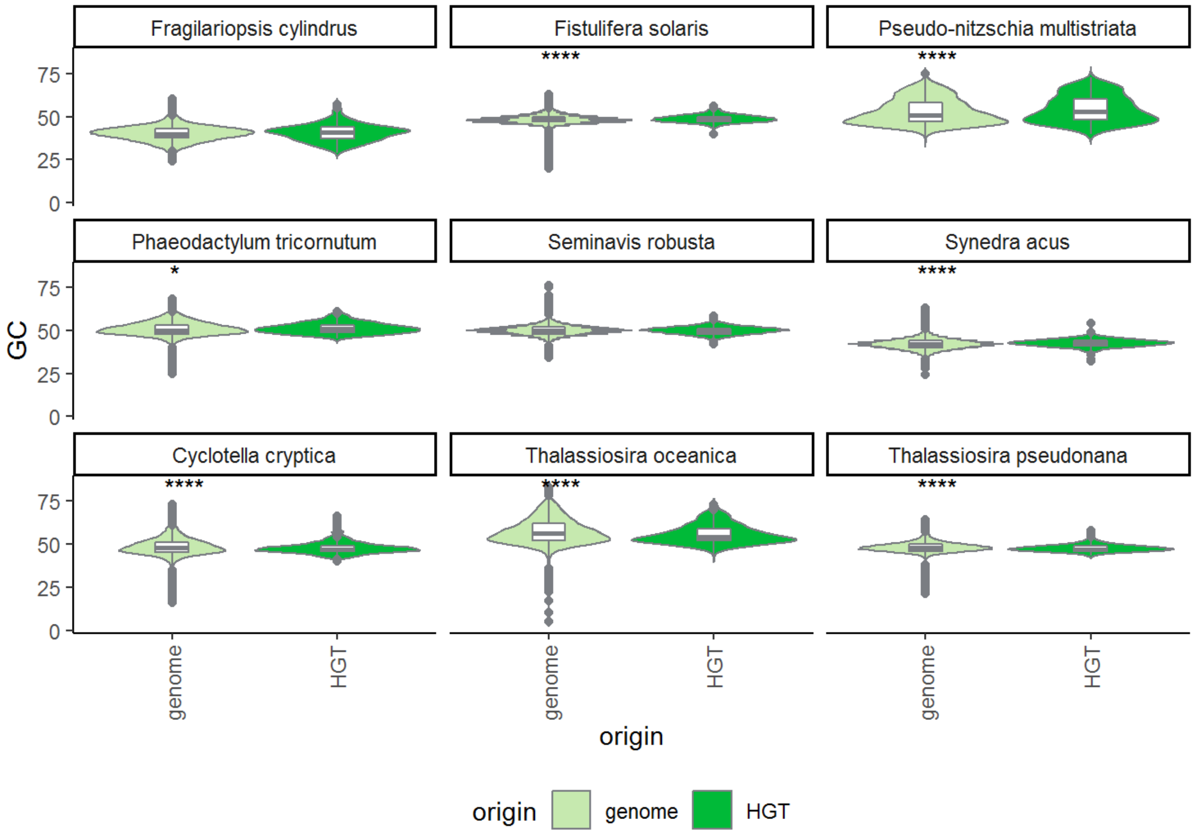

**Figure S1: GC distribution per origin for all nine diatom species.** The asterisks denote a statistical difference (Wilcoxon rank sum test) per type within the same age category and have the following confidence range for p-values; * : ≤ 0.05, ** : ≤ 0.01, *** : ≤ 0.001, **** : 0.0001.

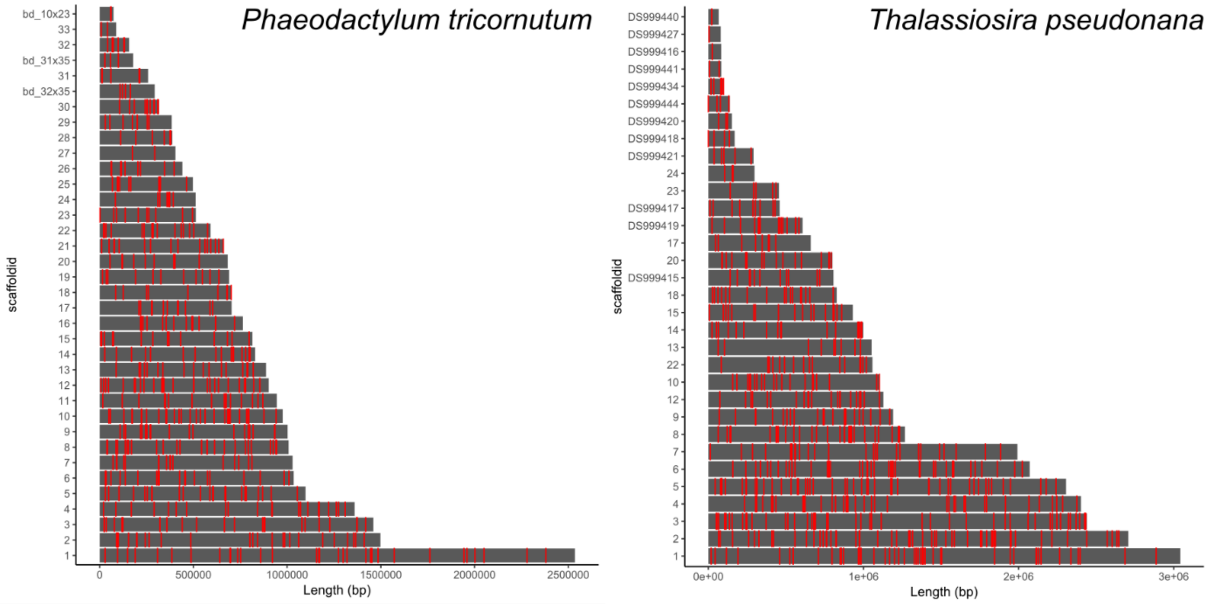

**Figure S2: Distribution of HGT genes across chromosome-level diatom genomes.** Distribution of HGT genes across the genome of *Phaeodactylum tricornutum* (left) and *Thalassiosira pseudonana* (right).

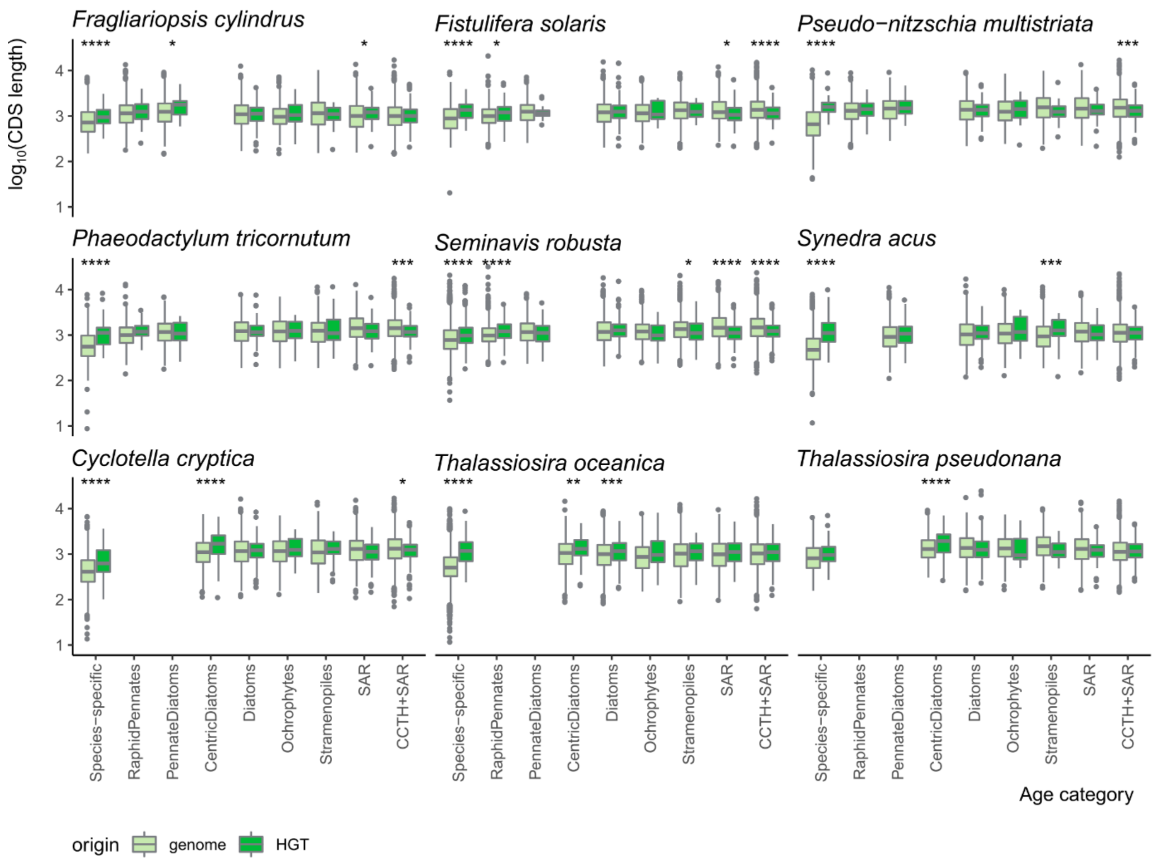

**Figure S3: CDS length per age category per origin across species.** CDS lengths were log10 transformed. The asterisks denote a statistical difference (Wilcoxon rank sum test) per type within the same age category and have the following confidence range for p-values; * : ≤ 0.05, ** : ≤ 0.01, *** : ≤ 0.001, **** : 0.0001.

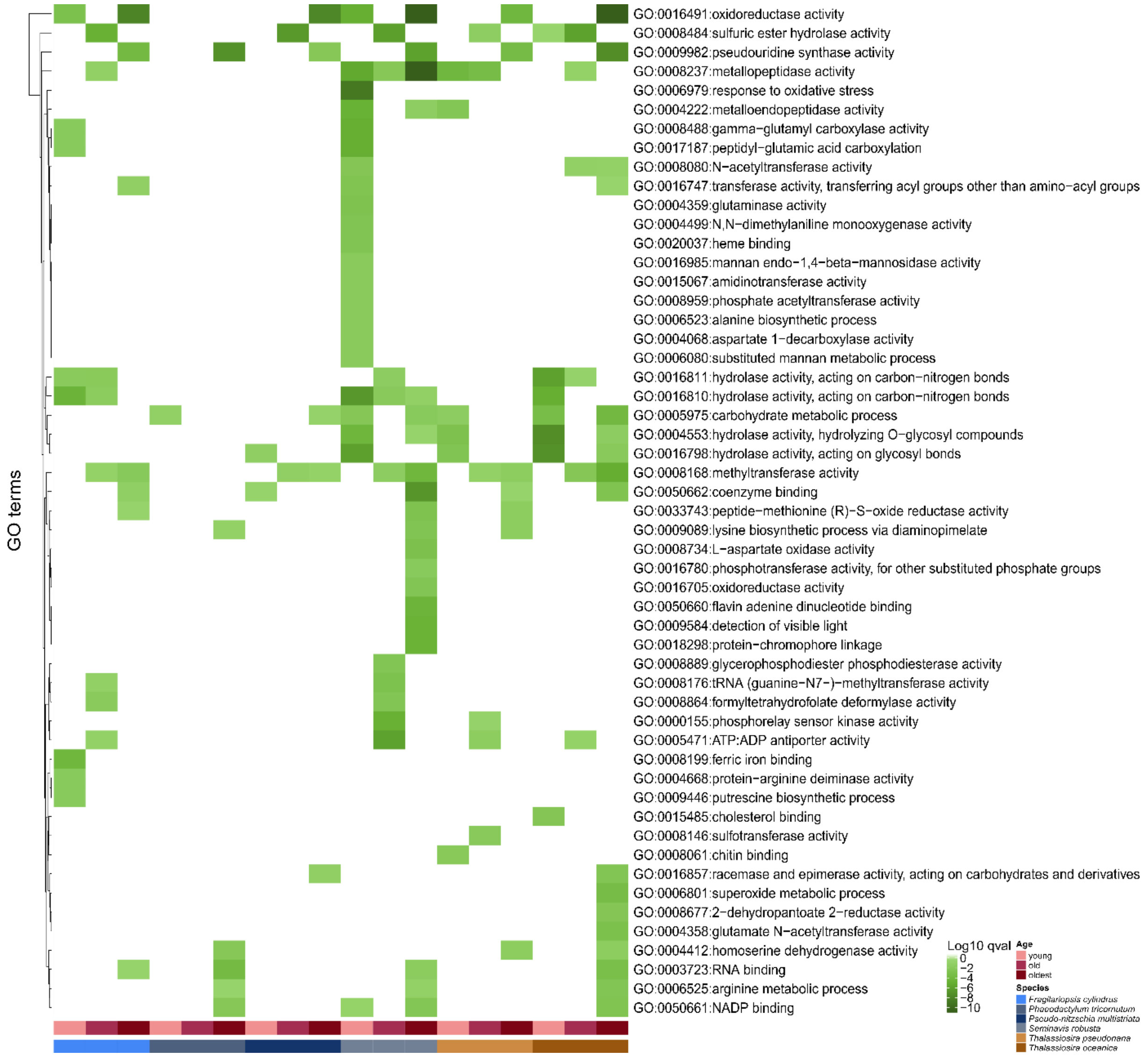

**Figure S4: Gene ontology enrichment of HGT genes across diatoms.** Only GO terms having an enrichment of at least ≤5x10$^{-3}$ are shown.

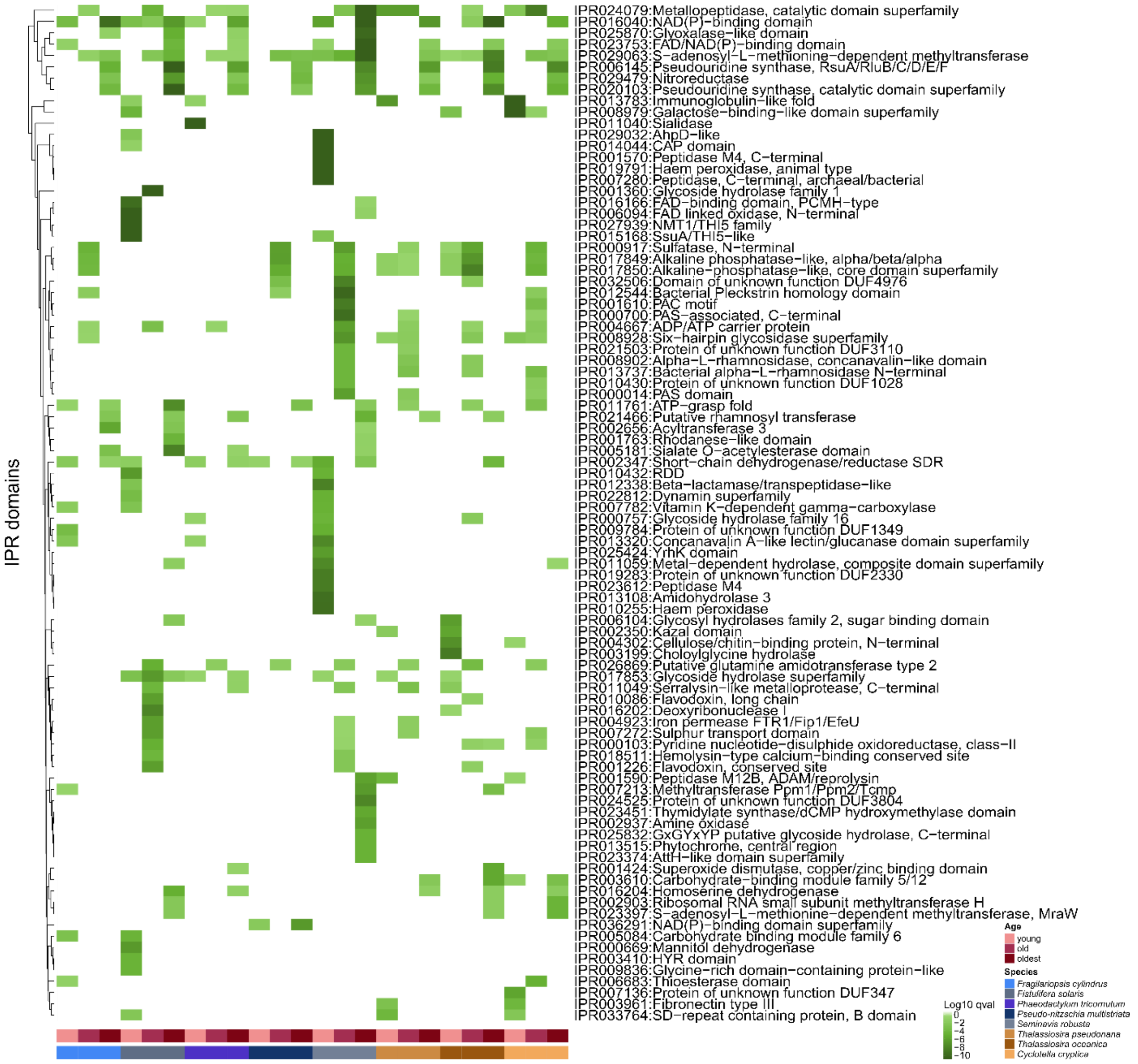

**Figure S5: Functional domain enrichment of HGT genes across diatoms.** Only Interpro domains having an enrichment of at least ≤5x10$^{-5}$ are shown.

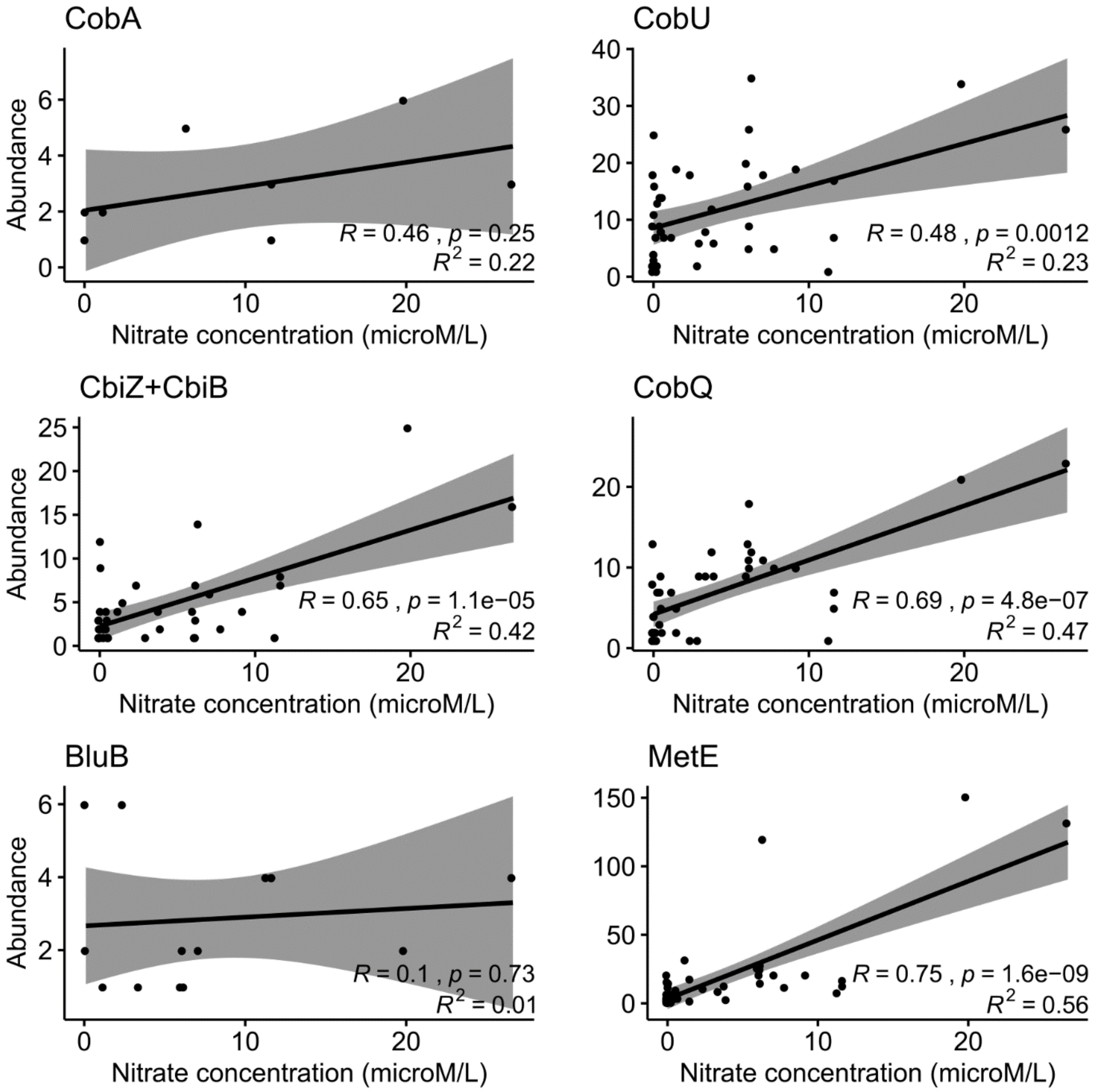

**Figure S6: Correlation between diatom gene abundance and nitrate concentration at surface depth.** The cobalamin-independent enzyme MetE is included as a reference.

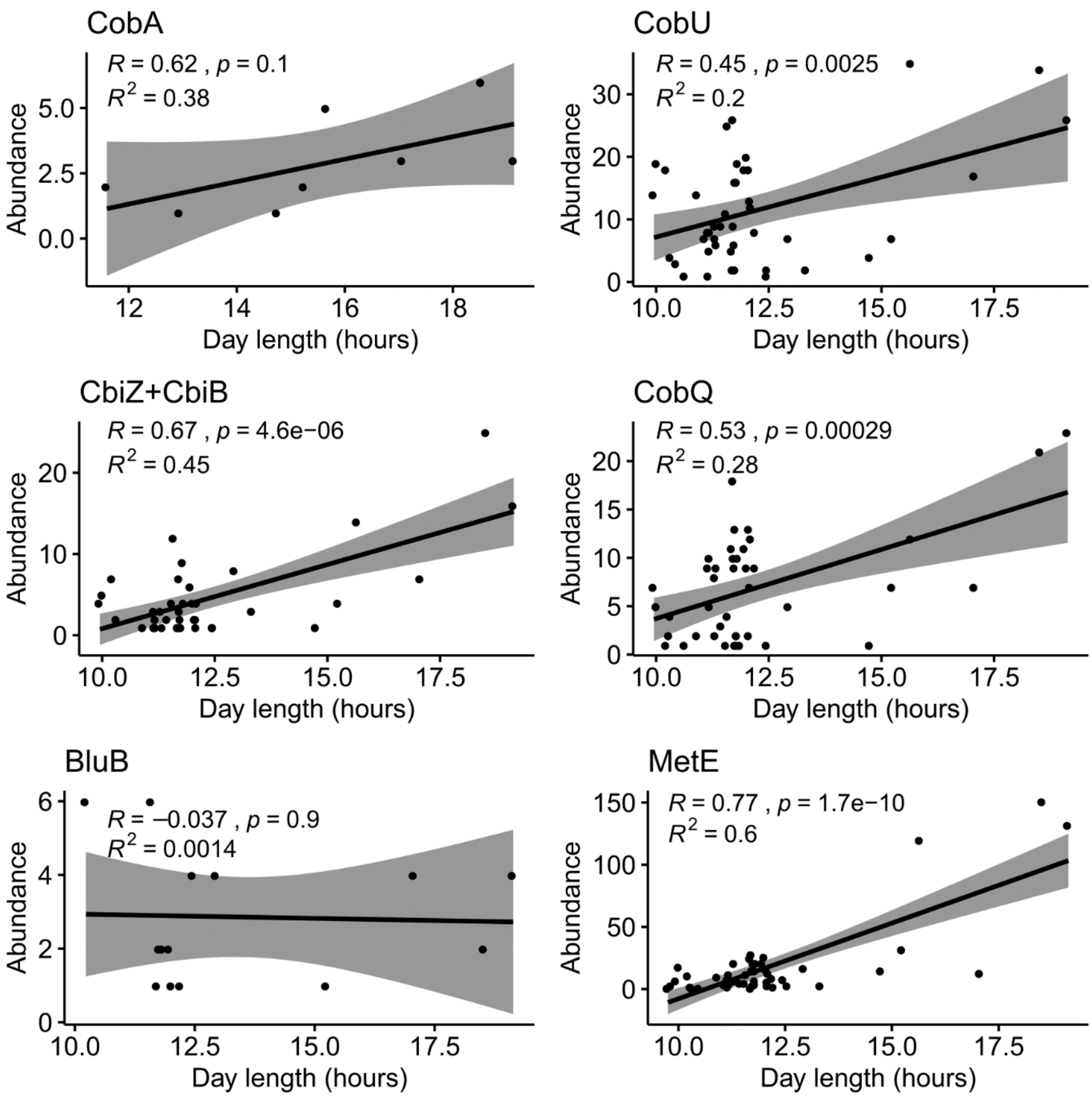

**Figure S7: Correlation between diatom gene abundance and sampling day length at surface depth.** The cobalamin-independent enzyme MetE is included as a reference.

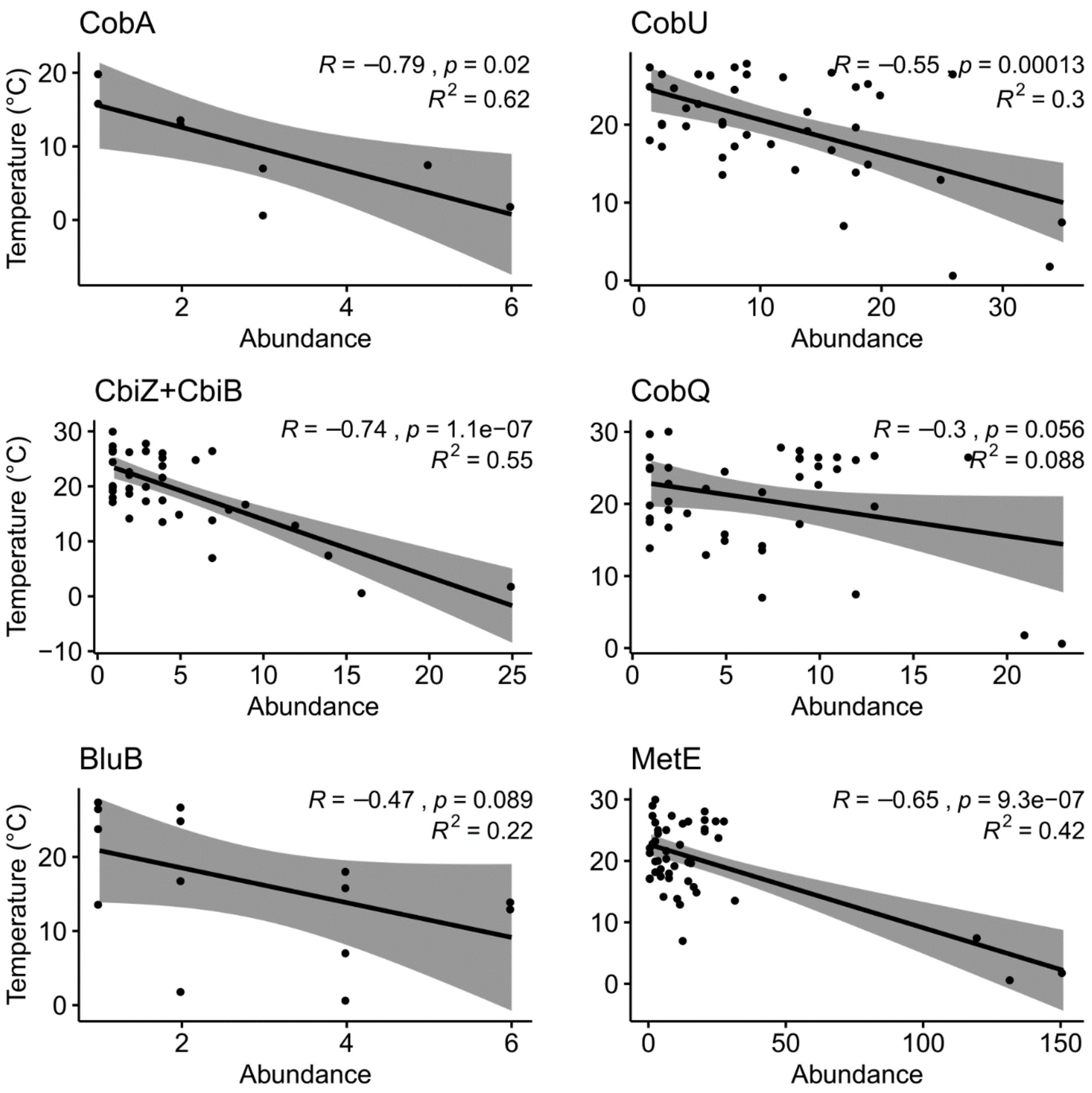

**Figure S8: Correlation between diatom gene abundance and water temperature at surface depth.** The cobalamin-independent enzyme MetE is included as a reference.

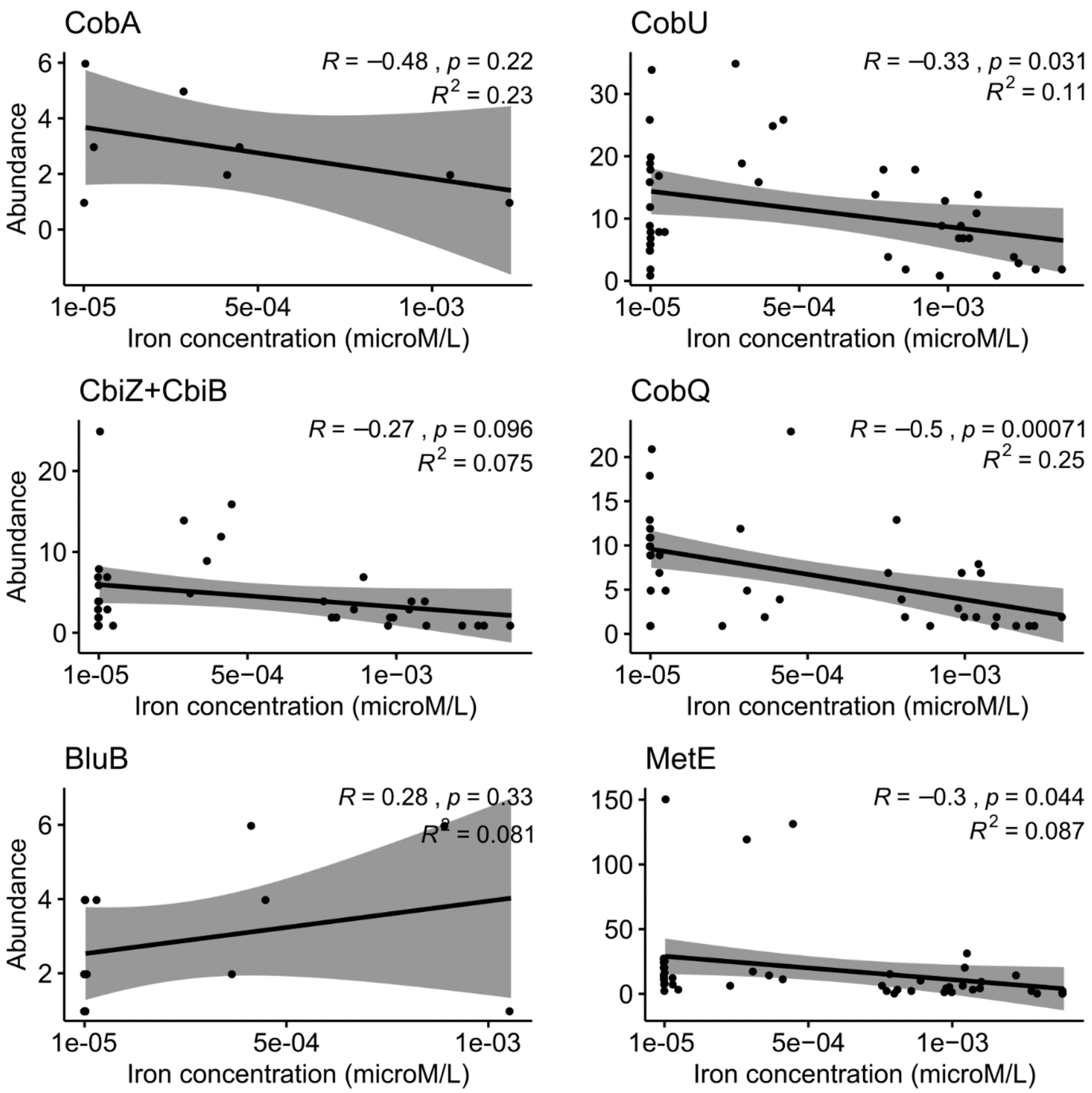

**Figure S9: Correlation between diatom gene abundance and iron concentration at surface depth.** The cobalamin-independent enzyme MetE is included as a reference.

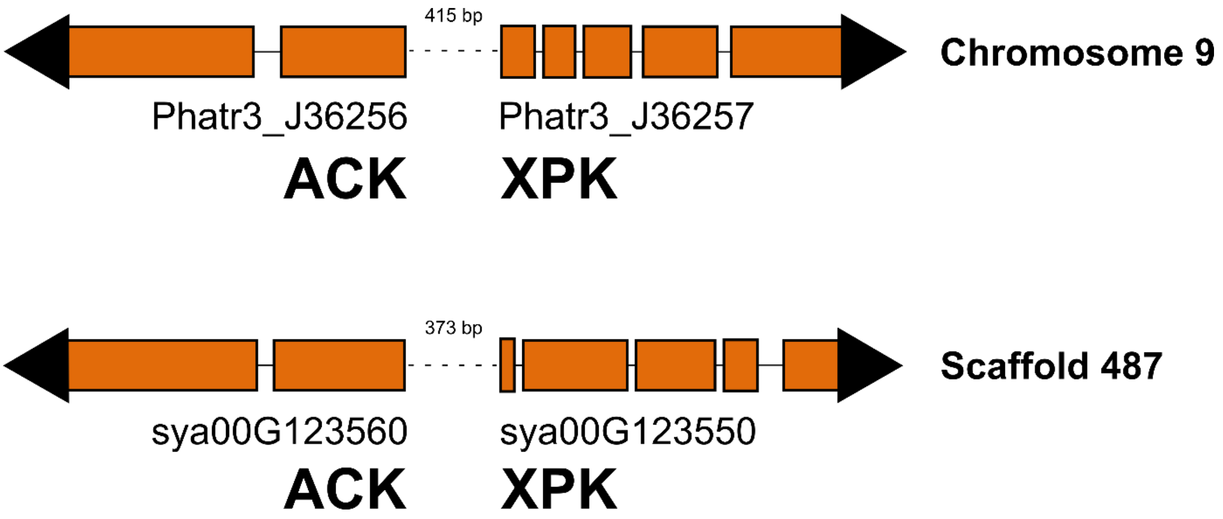

**Figure S10: Gene organization of the bifid shunt operon.** Syntenic organization of ACK and XPK in *Phaeodactyum tricornutum* (above) and *Synedra acus* (below). Exons are indicated by orange blocks, introns by solid lines and the intergenic region by a dashed line. The length of the intergenic region is displayed in number of basepairs. Direction of transcription is shown by the placement of an arrow.

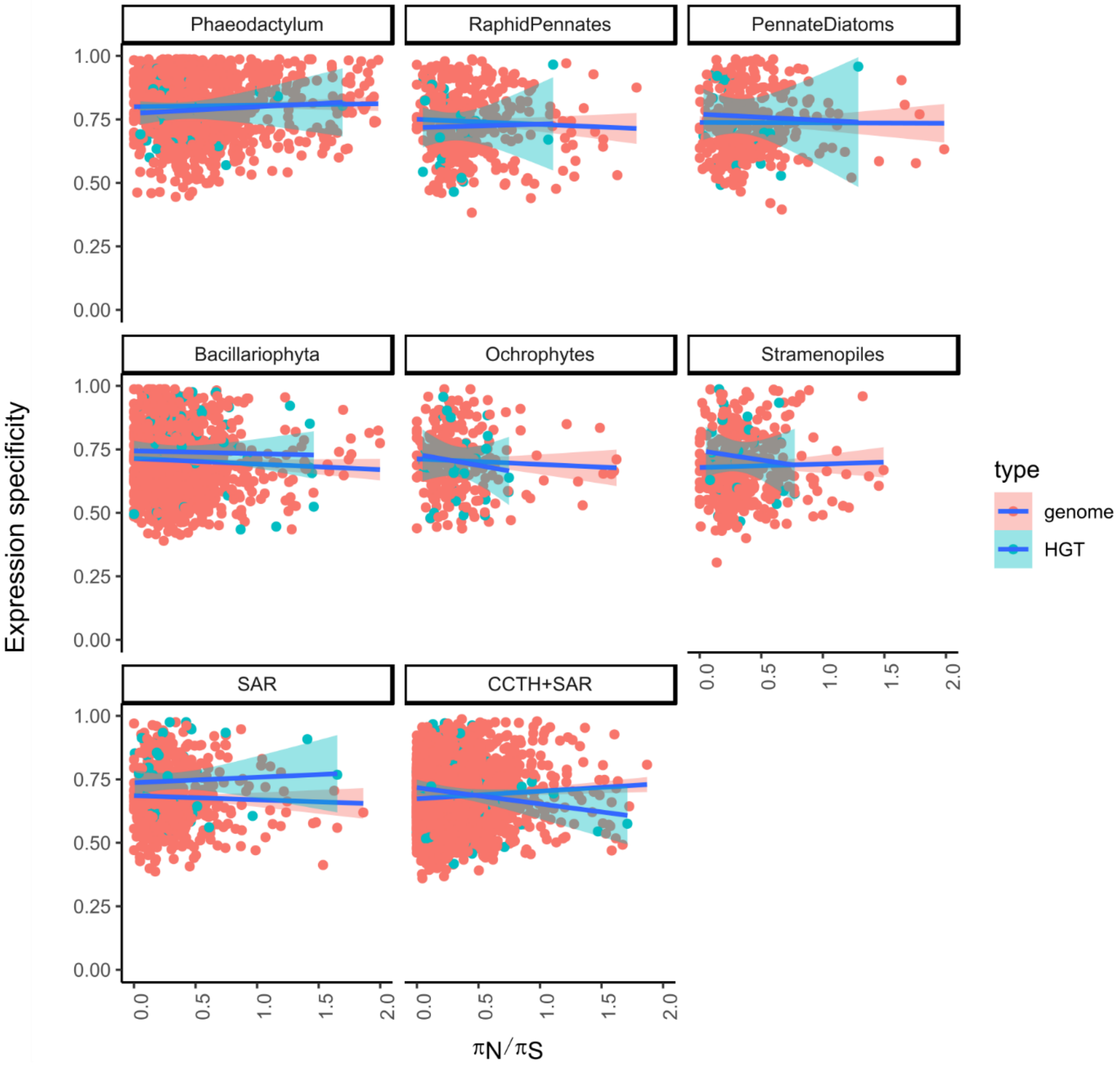

**Figure S11: Correlation between expression specificity and selection pressure.** Comparison of expression specificity and selection pressure across age categories and origin in *P. tricornutum*.

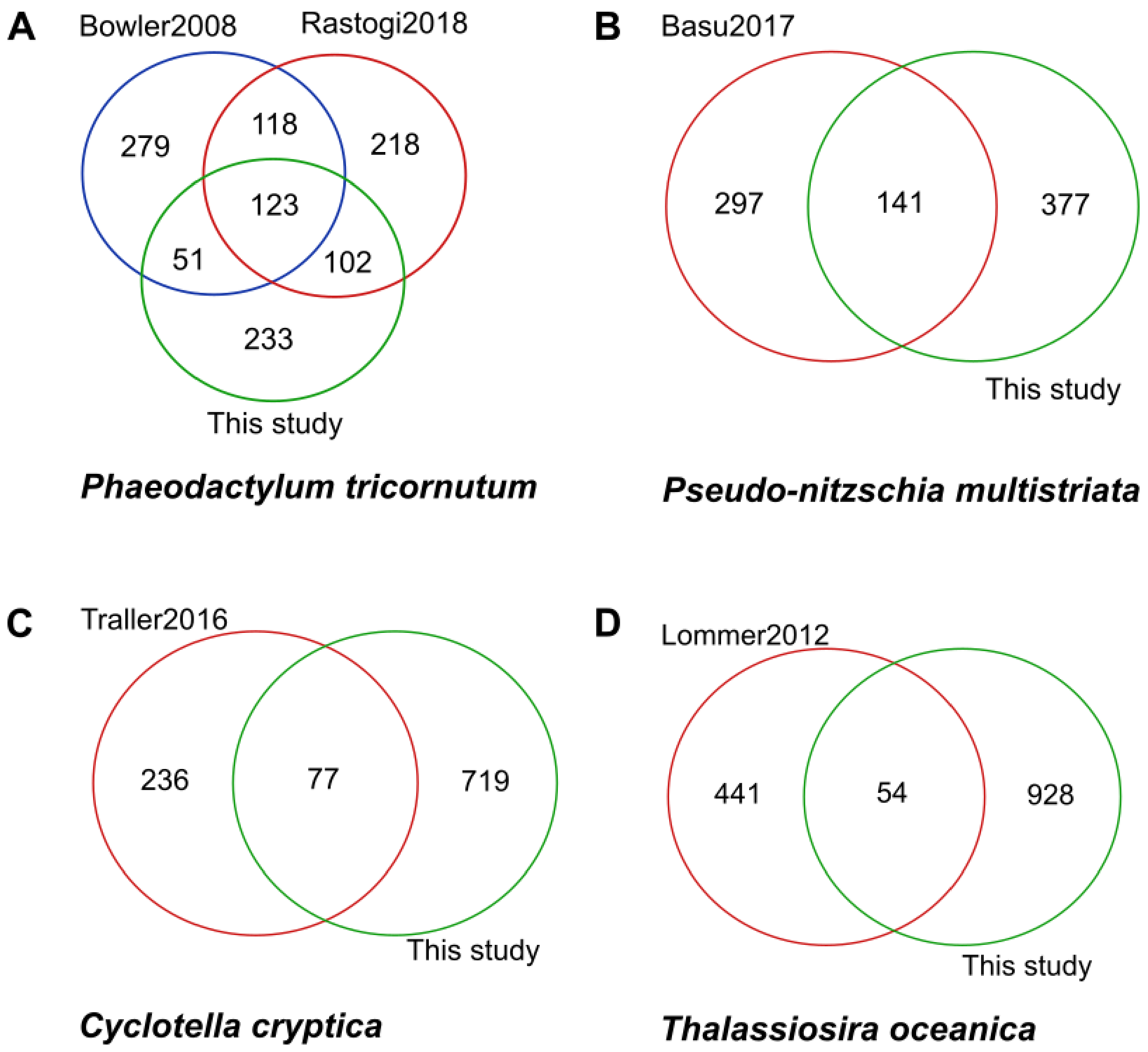

**Figure S12: Comparison between different published HGT sets and this study.** Comparison between different published HGT sets and the data set described in this study for *Phaeodactylum tricornutum* (A), *Pseudo-nitzschia multistriata* (B), *Cyclotella cryptica* (C) and *Thalassiosira oceanica* (D).

### Supplementary Tables

**Table S1: Overview of genomes used in this study.**

| Species | Lineage | PubmedID |
|---|---|---|
| *Blastocystis hominis* | Opalozoa, Stramenopiles | 21439036 |
| *Aplanochytrium kerguelense* | Labyrinthulea, Stramenopiles | JGI |
| *Aurantiochytrium limacinum* | Labyrinthulea, Stramenopiles | JGI |
| *Schizochytrium aggregatum* | Labyrinthulea, Stramenopiles | JGI |
| *Hyphochytrium catenoides* | Hyphochytriomycetes, Stramenopiles | 29321239 |
| *Pythium ultimum* | Oomycetes, Stramenopiles | 20626842 |
| *Ectocarpus siliculosus* | Phaeophycaea, Stramenopiles | 27870061 |
| *Nannochloropsis gaditana* | Eustigmatophyceae, Stramenopiles | 23966634 |
| *Thalassiosira oceanica* | Polar centrics, Diatoms, Stramenopiles | 22835381 |
| *Thalassiosira pseudonana* | Polar centrics, Diatoms, Stramenopiles | 15459382 |
| *Cyclotella cryptica* | Polar centrics, Diatoms, Stramenopiles | 27933100 |
| *Synedra acus* | Araphid pennates, Diatoms, Stramenopiles | 25937221 |
| *Seminavis robusta* | Raphid pennates, Diatoms, Stramenopiles | *in house* |
| *Phaeodactylum tricornutum* | Raphid pennates, Diatoms, Stramenopiles | 29556065 |
| *Fragilariopsis cylindrus* | Raphid pennates, Diatoms, Stramenopiles | 28092920 |
| *Fistulifera solaris* | Raphid pennates, Diatoms, Stramenopiles | 25634988 |
| *Pseudo-nitzschia multistriata* | Raphid pennates, Diatoms, Stramenopiles | 28429538 |
| *Paramecium tetraurelia* | Alveolata | 17086204 |
| *Bigelowiella natans* | Rhizaria | 16760254 |
| *Emiliania huxleyi* | Haptophyceae | 23760476 |

**Table S2: Expanded HGT gene families.** The following abbreviations for diatoms were used; tho: *T. oceanica*, tps: *T. pseudonana*, cycr: *C. cryptica*, sac: *S. acus*, sro: *S. robusta*, ptri: *P. tricornutum*, fcy: *F. cylindrus*, pmu: *P. multistriata*. All phylogenetic trees can be looked up by their gene family identifier (ORTHO01HGTXXXXXX) and are available in the supplemental dataset on Zenodo.

| Gene family | Function | Expansion | tho | cycr | tps | sac | sro | ptri | fcy | pmu |
|---|---|---|---|---|---|---|---|---|---|---|
| 000077 | DNA integration | tho | 6 | 0 | 0 | 0 | 1 | 0 | 2 | 0 |
| 000231 | Sulfatase | sro | 6 | 5 | 2 | 4 | 7 | 1 | 4 | 6 |
| 000322 | PAS domain | sro | 2 | 5 | 5 | 3 | 9 | 0 | 0 | 0 |
| 000364 | Peptidase M6-like | sac,sro | 0 | 1 | 0 | 6 | 4 | 0 | 1 | 0 |
| 000370 | Flavin monooxygenase FMO | fcy,sro | 0 | 0 | 0 | 0 | 5 | 0 | 2 | 1 |
| 000399 | Metallopeptidase | sac, tps | 0 | 2 | 4 | 5 | 0 | 0 | 0 | 0 |
| 000408 | ATP-grasp fold | fcy, pmu, sac | 0 | 3 | 3 | 5 | 3 | 2 | 6 | 5 |
| 000416 | Multicopper oxidase | sac | 0 | 0 | 0 | 5 | 2 | 0 | 1 | 1 |
| 000454 | Peptide methionine sulphoxide reductase MsrB | sro | 0 | 2 | 2 | 4 | 5 | 2 | 4 | 3 |
| 000518 | P-loop containing nucleoside triphosphate hydrolase | sro, ptri | 6 | 6 | 5 | 6 | 14 | 13 | 4 | 3 |
| 000700 | Metallo-dependent phosphatase-like | sac | 2 | 2 | 4 | 16 | 2 | 1 | 5 | 5 |
| 000729 | PDZ domain | sro | 0 | 3 | 2 | 0 | 31 | 0 | 3 | 2 |
| 000823 | Glycoside hydrolase | sro,tho | 4 | 2 | 1 | 0 | 6 | 2 | 1 | 0 |
| 001011 | Metallopeptidase | sro | 1 | 0 | 0 | 0 | 13 | 0 | 0 | 0 |
| 001143 | Phytase-like domain | tho | 7 | 1 | 1 | 0 | 1 | 1 | 2 | 2 |
| 001185 | Spondin, N-terminal | sro | 2 | 2 | 2 | 0 | 6 | 2 | 0 | 0 |
| 001894 | Unknown | sac | 0 | 0 | 0 | 5 | 1 | 0 | 0 | 0 |
| 002025 | Histidine kinase | sro | 3 | 4 | 2 | 2 | 8 | 1 | 0 | 0 |
| 002026 | Carbohydrate-binding module family 5/12 | cycr,tho | 16 | 7 | 3 | 0 | 0 | 0 | 0 | 0 |
| 002179 | Alpha-L-rhamnosidase | cycr,tho | 5 | 4 | 3 | 0 | 3 | 0 | 1 | 0 |
| 002202 | Superoxide dismutase, copper/zinc binding domain | sac,tho | 4 | 0 | 0 | 4 | 0 | 2 | 1 | 1 |
| 002203 | Amine oxidase | sac,sro | 2 | 0 | 0 | 5 | 12 | 1 | 0 | 0 |
| 002376 | CAP domain | sac,sro | 0 | 0 | 0 | 8 | 11 | 1 | 2 | 2 |
| 002599 | Unknown | tho | 20 | 0 | 0 | 0 | 1 | 0 | 0 | 0 |
| 002816 | Alpha/Beta hydrolase fold | ptri,sro | 3 | 0 | 1 | 3 | 6 | 5 | 2 | 1 |
| 003267 | Aminotransferase + Tubulin-tyrosine ligase | tps | 1 | 2 | 6 | 1 | 1 | 1 | 1 | 1 |
| 003534 | DPH far-red/red light sensor | sac,sro | 0 | 1 | 1 | 7 | 4 | 1 | 0 | 0 |
| 003583 | Unknown | cycr,tps | 0 | 7 | 6 | 1 | 1 | 1 | 1 | 1 |
| 003977 | S-adenosylmethionine: tRNA-ribosyltransferase-isomerase | sac | 2 | 0 | 0 | 5 | 2 | 0 | 0 | 1 |
| 004024 | P-loop containing nucleoside triphosphate hydrolase | sac,sro | 0 | 0 | 0 | 3 | 4 | 0 | 2 | 1 |
| 004133 | Unknown | tho,tps | 5 | 1 | 12 | 0 | 0 | 0 | 0 | 0 |
| 004304 | Tetratricopeptide repeat | cycr, tho,tps | 4 | 3 | 4 | 0 | 0 | 0 | 0 | 0 |
| 004743 | Prolyl 4-hydroxylase | tho | 9 | 1 | 0 | 0 | 0 | 0 | 0 | 0 |
| 004768 | GTP-binding domain | sro | 0 | 0 | 0 | 0 | 9 | 0 | 1 | 1 |
| 004770 | Glyoxalase-like domain | sro | 0 | 0 | 0 | 0 | 8 | 2 | 1 | 1 |
| 005013 | Unknown | cycr, tho,tps | 3 | 6 | 3 | 1 | 0 | 0 | 0 | 0 |

| | | | | | | | | | | |
|---|---|---|---|---|---|---|---|---|---|---|
| 005077 | ATP-grasp fold | cycr, tho,tps | 5 | 3 | 3 | 1 | 1 | 0 | 1 | 1 |
| 005106 | Galactose-binding-like domain superfamily | fcy, ptri | 0 | 1 | 0 | 0 | 1 | 5 | 4 | 2 |
| 005155 | Protein of unknown function DUF1349 | fcy,sro | 0 | 0 | 0 | 0 | 5 | 1 | 2 | 0 |
| 005382 | PH-like domain superfamily | sro | 1 | 2 | 1 | 0 | 6 | 0 | 2 | 2 |
| 005501 | P-loop containing nucleoside triphosphate hydrolase | sro | 0 | 0 | 0 | 0 | 6 | 1 | 2 | 2 |
| 005514 | Holliday junction resolvase RusA-like | ptri,sro | 1 | 0 | 1 | 1 | 5 | 3 | 1 | 0 |
| 005775 | Nucleophile aminohydrolases | cycr | 1 | 8 | 1 | 0 | 3 | 0 | 0 | 0 |
| 005825 | Cytochrome P450 | sro | 0 | 1 | 0 | 0 | 10 | 0 | 0 | 1 |
| 005865 | Unknown | tps | 0 | 2 | 7 | 0 | 0 | 0 | 1 | 1 |
| 006462 | Metallopeptidase | sro | 0 | 0 | 0 | 0 | 7 | 2 | 0 | 0 |
| 006467 | Unknown | sac | 0 | 0 | 0 | 11 | 1 | 0 | 0 | 0 |
| 006701 | Metallopeptidase | cycr, tho,tps | 4 | 3 | 3 | 0 | 1 | 0 | 0 | 0 |
| 006767 | Pectin lyase fold | fcy | 1 | 2 | 0 | 1 | 0 | 0 | 7 | 0 |
| 006838 | Unknown | sac,tho | 4 | 1 | 1 | 5 | 0 | 0 | 0 | 0 |
| 006877 | Ketopantoate reductase | tho | 5 | 1 | 0 | 1 | 1 | 1 | 1 | 0 |
| 006980 | Unknown | ptri | 0 | 0 | 0 | 0 | 1 | 7 | 0 | 0 |
| 006994 | YrhK domain | sro | 0 | 0 | 0 | 0 | 5 | 1 | 1 | 0 |
| 007014 | Unknown | sac | 0 | 1 | 0 | 9 | 0 | 0 | 0 | 0 |
| 007214 | Unknown | cycr | 0 | 8 | 2 | 0 | 0 | 0 | 0 | 0 |
| 007566 | von Willebrand factor, type A | sac | 0 | 0 | 0 | 7 | 0 | 0 | 2 | 1 |
| 007599 | Unknown | ptri,sac | 0 | 0 | 0 | 6 | 1 | 3 | 0 | 0 |
| 007931 | Unknown | cycr, tho | 5 | 3 | 1 | 0 | 0 | 0 | 0 | 0 |
| 007964 | Sulfotransferase | tps | 1 | 2 | 6 | 0 | 0 | 0 | 0 | 0 |
| 008041 | Peptidase M66 domain | sac | 2 | 1 | 0 | 6 | 0 | 0 | 0 | 0 |
| 008138 | Protein of unknown function DUF3804 | sro | 1 | 0 | 0 | 0 | 6 | 0 | 0 | 0 |
| 008163 | Bulb-type lectin domain | fcy, pmu, sac | 0 | 0 | 0 | 2 | 0 | 0 | 4 | 3 |
| 008164 | Ice-binding proteins | fcy | 2 | 0 | 0 | 0 | 0 | 0 | 6 | 0 |
| 008818 | Tetratricopeptide repeat | tho | 7 | 0 | 1 | 0 | 0 | 0 | 0 | 0 |
| 009138 | Metallo-dependent phosphatase-like | cycr | 0 | 6 | 1 | 0 | 0 | 0 | 0 | 0 |
| 009256 | Phosphoglycerate/ bisphosphoglycerate mutase | tho | 5 | 1 | 1 | 0 | 0 | 0 | 0 | 0 |
| 009470 | XPK | sac | 0 | 0 | 0 | 5 | 0 | 1 | 0 | 0 |
| 009493 | Unknown | sac | 1 | 0 | 0 | 6 | 0 | 0 | 0 | 0 |

**Table S3: Overview of all discussed HGT gene families.** The copy number is give per species. he following abbreviations for diatoms were used; tho: *T. oceanica*, tps: *T. pseudonana*, cycr: *C. cryptica*, sac: *S. acus*, sro: *S. robusta*, ptri: *P. tricornutum*, fcy: *F. cylindrus*, pmu: *P. multistriata*. The last column indicates whether this gene was already confirmed by another study to be horizontally transferred. All phylogenetic trees can be looked up by their gene family identifier (ORTHO01HGTXXXXXX) and are available in the supplemental dataset on Zenodo.

| | | | **tho** | **cycr** | **tps** | **sac** | **sro** | **ptri** | **fso** | **fcy** | **pmu** | **Ref** |
|---|---|---|---|---|---|---|---|---|---|---|---|---|
| Iron | FBP1 | 006985 | 1 | 0 | 0 | 0 | 1 | 1 | 5 | 2 | 1 | [1] |
| | Iron permease | 006316 | 0 | 1 | 3 | 2 | 1 | 0 | 4 | 1 | 0 | / |
| | FTN | 011114 | 0 | 0 | 0 | 0 | 0 | 0 | 0 | 1 | 0 | [2] |
| | Proteorhodopsin | 067862 | 0 | 0 | 0 | 0 | 0 | 0 | 0 | 1 | 0 | [3] |
| | Proteorhodopsin | 067193 | 0 | 0 | 0 | 0 | 0 | 0 | 0 | 1 | 0 | [3] |
| Vitamin B12 | CobN | 017541 | 0 | 0 | 0 | 0 | 1 | 0 | 0 | 1 | 0 | / |
| | CobA/CobO | 068467 | 0 | 0 | 0 | 0 | 0 | 0 | 0 | 1 | 0 | / |
| | CobQ/CbiP | 010641 | 0 | 0 | 1 | 0 | 0 | 0 | 0 | 1 | 2 | / |
| | CbiZ+CbiB | 008486 | 0 | 3 | 0 | 0 | 1 | 0 | 0 | 1 | 1 | / |
| | CobU/CobP | 009121 | 2 | 1 | 0 | 0 | 1 | 0 | 0 | 1 | 1 | [4,5] |
| | BluB | 013423 | 0 | 0 | 0 | 0 | 0 | 0 | 0 | 1 | 1 | [5] |
| Micro-nutrients | ThiD+ThiE | 001873 | 1 | 1 | 2 | 1 | 2 | 1 | 0 | 2 | 1 | [6] |
| | Thi5-like | 009317 | 1 | 0 | 1 | 0 | 0 | 1 | 0 | 1 | 0 | [6,7] |
| | MenA | 004876 | 1 | 1 | 1 | 1 | 1 | 1 | 2 | 1 | 1 | [5,6,8] |
| | MerC | 006910 | 1 | 1 | 0 | 1 | 1 | 1 | 2 | 1 | 1 | / |
| Sensing | DPH1 | 003534 | 0 | 1 | 1 | 7 | 4 | 1 | 0 | 0 | 0 | [7] |
| | Ice-binding-like | 008164 | 0 | 0 | 0 | 0 | 0 | 0 | 0 | 6 | 0 | / |
| Cell wall | CDA | 004440 | 2 | 1 | 1 | 2 | 1 | 1 | 2 | 1 | 1 | [6,7,9] |
| | Rhamnosyl transferase | 001567 | 4 | 4 | 3 | 4 | 4 | 3 | 3 | 3 | 2 | / |
| | TupA-like | 005583 | 1 | 3 | 0 | 3 | 0 | 1 | 0 | 0 | 0 | [6] |
| Carbon metabolism | Carbamate kinase | 006183 | 1 | 1 | 1 | 1 | 2 | 1 | 2 | 1 | 1 | [5–7,10] |
| | Ornithine cyclodeaminase | 007884 | 1 | 1 | 1 | 1 | 1 | 1 | 0 | 1 | 1 | [6,7,10] |
| | Allantoin synthase | 003479 | 1 | 2 | 1 | 1 | 1 | 1 | 2 | 1 | 1 | [6,7,11] |
| | Phosphate acetyltransferase | 001020 | 2 | 2 | 2 | 2 | 3 | 2 | 4 | 1 | 1 | [5–7] |
| | ACK | 005298 | 0 | 0 | 0 | 1 | 1 | 1 | 0 | 0 | 0 | [6,7,12] |
| | XPK | 009470 | 0 | 0 | 0 | 5 | 0 | 1 | 0 | 0 | 0 | [6,7,12] |
| | Phosphofructokinase | 004987 | 2 | 3 | 1 | 0 | 2 | 1 | 2 | 1 | 1 | [5–7] |
| | Fba4 | 006382 | 1 | 1 | 1 | 1 | 1 | 1 | 2 | 1 | 2 | [5–7,13] |
| | Phosphopentose epimerase | 000787 | 4 | 3 | 2 | 1 | 1 | 1 | 2 | 2 | 3 | [5–7,14] |
| | D-lactate dehydrogenase | 005413 | 1 | 1 | 1 | 4 | 1 | 1 | 2 | 1 | 1 | [5,7] |
| | Pyruvate kinase-like | 009232 | 1 | 1 | 0 | 2 | 1 | 1 | 1 | 0 | 0 | [6] |
| | Xylanase | 005372 | 2 | 1 | 1 | 1 | 1 | 1 | 2 | 1 | 1 | [5–7] |
| | Glucanase | 000823 | 4 | 2 | 1 | 0 | 6 | 2 | 3 | 1 | 0 | [4,6,7] |
| | Glucosidase | 000466 | 0 | 0 | 0 | 2 | 2 | 1 | 2 | 1 | 1 | [5–7] |
| Amino acid metabolism | Asd | 001898 | 2 | 1 | 1 | 1 | 3 | 1 | 1 | 1 | 1 | [5–7] |
| | DapA | 001023 | 1 | 2 | 2 | 3 | 3 | 1 | 2 | 1 | 1 | [5–7] |
| | ThrA | 001521 | 2 | 2 | 2 | 3 | 2 | 2 | 4 | 2 | 1 | [5–7] |
| | Tryptophan synthase β chain | 001860 | 0 | 0 | 0 | 0 | 0 | 1 | 0 | 0 | 0 | [6,7,15] |
| | Alanine racemase | 006705 | 0 | 3 | 1 | 1 | 1 | 1 | 2 | 1 | 1 | / |
| | ArgJ | 003184 | 3 | 1 | 1 | 1 | 1 | 1 | 2 | 1 | 1 | [5–7] |
| | LeuRS2 | 002519 | 2 | 1 | 1 | 3 | 1 | 1 | 2 | 1 | 1 | [6,7] |
| | GlyRS2 | 003331 | 1 | 1 | 1 | 2 | 1 | 1 | 2 | 1 | 1 | [5–7] |
| | TyrRS2 | 002118 | 1 | 3 | 1 | 2 | 1 | 1 | 2 | 1 | 1 | [5–7] |
| Nucleotide import | NTT2 | 001110 | 2 | 2 | 2 | 1 | 1 | 1 | 2 | 1 | 0 | [16] |
| | NTT5/6 | 000793 | 2 | 3 | 3 | 3 | 5 | 2 | 4 | 2 | 1 | [6,16] |

**Table S4. Mapping and polymorphism statistics for ten resequencing strains.**

| Strain | Clade | Mapping percentage | Coverage | Missing genes | Number of heterozygous SNPs | Number of homozygous SNPs | Fraction of heterozygous SNPs | Total fraction of SNPs |
|---|---|---|---|---|---|---|---|---|
| Pt1 | A | 89.84% | 30 | 117 | 283,851 | 999 | 99.65% | 1.04% |
| Pt2 | A | 90.46% | 51 | 113 | 278,280 | 4,852 | 98.29% | 1.03% |
| Pt3 | A | 90.55% | 189 | 109 | 287,408 | 777 | 99.73% | 1.05% |
| Pt4 | B | 87.64% | 150 | 171 | 149,022 | 164,285 | 47.56% | 1.14% |
| Pt5 | C | 90.46% | 48 | 248 | 109,755 | 128,456 | 46.07% | 0.87% |
| Pt6 | D | 88.71% | 39 | 190 | 235,556 | 107,105 | 68.74% | 1.25% |
| Pt7 | D | 87.61% | 47 | 172 | 236,010 | 106,280 | 68.95% | 1.25% |
| Pt8 | D | 86.23% | 61 | 203 | 225,004 | 112,018 | 66.76% | 1.23% |
| Pt9 | A | 88.12% | 65 | 131 | 262,253 | 15,466 | 94.43% | 1.01% |
| Pt10 | C | 90.30% | 49 | 255 | 103,515 | 128,607 | 44.60% | 0.85% |

**Table S5. Overview of expression data used to create expression compendia.**

| **Species** | **Studies** | **Conditions** | **Samples** |
|---|---|---|---|
| *Phaeodactylum tricornutum* | ERP013403,SRP022147,SRP035546, SRP040703,SRP056249,SRP056740, SRP074144,SRP074517,SRP075327, SRP075821,SRP092213,SRP096318, SRP100419,SRP100930,SRP103881, SRP156408 | 76 | 211 |
| *Seminavis robusta* | ERP013194, SRP199371, *in house* | 58 | 167 |
| *Thalassiosira pseudonana* | SRP022147,SRP057269,SRP066751, SRP106713,SRP109670 | 42 | 123 |
| *Fragilariopsis cylindrus* | ERP016846,ERP104856, SRP022147 | 13 | 34 |

### Supplementary Notes

**Supplementary Note 1: Functional exploration of HGT genes**

**Micronutrient availability**

Diatoms are able to bloom both in iron-rich coastal areas and the iron-poor open ocean. Several gene transfers have occurred which may have facilitated its expansion in a low-iron environment. Iron uptake occurs by high-affinity ferric reductases, multi-copper oxidases and a Fe(III) permease (FTR)[2]. The putative ferrichrome-binding protein FBP1, which is part of an iron-responsive cluster and adjacent to ferric reductase (FRE2) in *P. tricornutum*[1], was suggested to be of horizontal descent[1]. This was here confirmed, while also detecting FBP1 in *F. cylindrus* (2 copies), *P. multistriata* (1 copy), *S. robusta* (1 copy), *F. solaris* (5 copies) and *T. oceanica* (1 copy). Both FBP1 and FRE2 are up-regulated in *P. tricornutum* during iron limitation[17] (Figure SN1). Moreover, iron permease receives iron from a multi-copper oxidase for translocation across the cell membrane and was present and perceived as laterally transferred in the centric diatoms *C. cryptica* (1 copy) and *T. pseudonana* (2 copies) and the raphid diatoms *F. cylindrus* (1 copy), *S. robusta* (1 copy), *S. acus* (2 copies) and the alloploid *F. solaris* (4 copies).

To safely store the iron that was taken up, *Pseudo-nitzschia* and *Fragilariopsis* use the iron concentrating protein, ferritin (FTN)[18]. An in-depth phylogenetic analysis of ferritin comprising transcriptome data from the MMETSP project, undertaken by Groussman et al.[2], detected ferritin in 32 diatoms which originates from horizontal gene transfer, while ferritin in *P. tricornutum* and one copy in *Nanofrustulum sp*. belonged to a different clade. In agreement with this, horizontal gene transfer was detected in this study for the ferritin gene family in *F. cylindrus*, but not for *P. tricornutum*.

Proteorhodopsin (PR) genes where detected as highly expressed under iron limitation, while having high similarity to bacteria[19]. It was suggested that due to lack of trace metals required in proteorhodopsins, they can supplement ATP generation as a light driven proton pump in low-iron environments[3]. This study confirms the bacterial origin of the PR-genes in *F. cylindrus* and *Pseudo-nitzschia granii*[3], next to brown algae, dinophytes and haptophytes.

Iron is intricately linked to vitamin B1 (thiamine) due to the presence of iron–sulphur (Fe-S) cluster-containing enzymes involved in its biosynthesis. Thiamine monophosphate biosynthesis from 4-amino-2-methyl-5-hydroxymethylpyrimidine monophosphate is performed in diatoms by a single bifunctional protein containing phosphomethylpyrimidine kinase (ThiD) and thiamine monophosphate synthase (ThiE) domains, similar to TH1 in A. thaliana[20]. However, this gene is shown here to be of horizontal descent and believed to have independently evolved this gene composition. Moreover, a gene family containing the THI5-like domain was predicted to be horizontally transferred, but cannot unambiguously be linked to thiamine biosynthesis.

Another micronutrient in which HGT played an important role is menaquinone (vitamin K2). The first enzyme in its biosynthesis, MenA is shown to be HGT. The rest of the pathway is part of a single nuclear-encoded, composite gene and was proposed to have been transferred via EGT[8].

Finally, also the mercury transport protein MerC was found to be horizontally transferred. Interestingly, diatoms have been reported to be an imported vector in mercury sequestration[21].

**Cell wall components**

Diatoms produce a porous silica cell wall as shelter from the environment, called the frustule. A compound embedded within the frustule is chitin, a structural polysaccharide that contributes to the rigidity of the cells[22]. Although diatoms contain genes involved in chitin synthesis, only a few genera, such as *Thalassiosira* and *Cyclotella* are known to produce chitin[23]. Chitosan is a partially de-acetylated chitin derivative formed by chitin deacetylase (CDA)[9] and could also be important for cell wall integrity. Despite that Shao et al.[9] showed that CDA in centric diatoms is transferred from fungi and in pennates from proteobacteria, a single bacterial event was detected in this analysis. Moreover, a family containing a putative rhamnosyl transferase domain was detected, that could be of importance in the synthesis of surface polysaccharides. Finally, also a TupA-like ATPgrasp protein was detected in diatoms that could be involved in the biosynthesis of cell surface polysaccharides. Interestingly, a member of this gene family in *P. tricornutum* (Phatr3_J47780) displayed nitrate-specific expression[24]. A proteomic analysis in *T. pseudonana* revealed an upregulation of extracellular polysaccharides production during nitrogen depletion, as part of the nutrient stress response[25].

**Nucleotide transport**

In diatoms, nucleotide metabolism occurs in the cytosol and nucleotide transporters (NTTs) are required for their transport to the plastid[26]. Analogous to[16,26], NTT1 which acts as a proton-dependent adenine nucleotide importer was not identified as HGT, while *P. tricornutum* NTT2 and NTT5/NTT6 and its orthologs were acquired by HGT in two separate events.

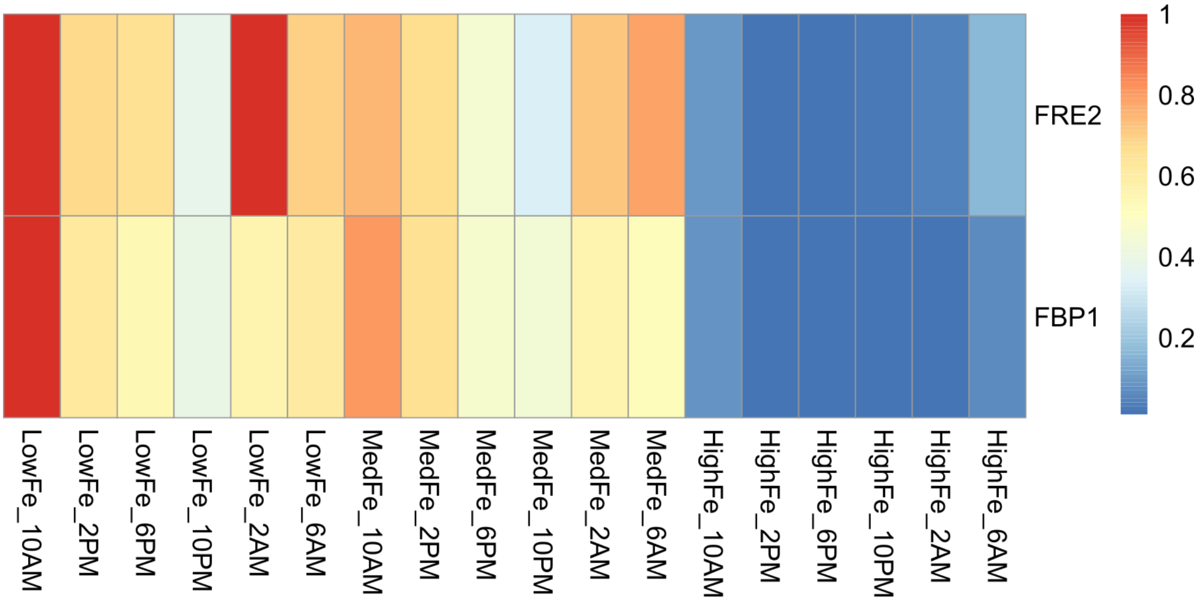

**Figure SN1: Expression of an iron-responsive cluster in fluctuating iron concentrations during the diel cycle.** Expression of the iron-responsive cluster FRE2/FBP1 at low, intermediate, and high levels of dissolved Fe during the diel cycle.

**Supplementary Note 2: Horizontal gene retention across different *P. tricornutum* strains**

Conservation of species-specific HGT genes across different strains can confirm their horizontally derived origin rather than point to contamination. The fraction of mapped reads varied from 86.23% to 90.55%, with a mean coverage ranging from 30 to 189X (Table S4). This allowed for a careful examination of the absence pattern of genes per strain. Only 256 genes of the whole proteome were categorized as belonging to the dispensable gene set of *P. tricornutum* and 101 were unique to the reference strain, in contrast to 11,821 genes (97%) of genes which are retained across all ten strains. The number of missing genes per individual strain ranged from 109 to 255 genes (Table S4). Nineteen HGT candidates in *P. tricornutum* were lost in at least one strain, which is 4% of all detected HGT genes (Figure SN2) (Table SN1). These ten strains could be divided into four genetic clades (A-D), with the reference belonging to clade A[27]. While in clade A at most one dispensable HGT gene is lost, this increases to half of these genes in strain Pt5 (clade C). For example, Phatr3_J16982, a glutathione S-transferase, was only missing in Pt5, while the pseudouridine synthase Phatr3_J46957 was absent from clade C and D (Pt6,7,8,5,10). The SDR oxidoreductase Phatr3_J5780 on the other hand, was absent in all members of clade D (Pt6,7,8).

**Table SN1: Missing HGT genes across ten *P. tricornutum* resequencing strains.**

| Gene ID | Family | Age | Missing strain | Missing clade |
|---|---|---|---|---|
| Phatr3_J44991 | ORTHO01HGT002816 | Bacillariophyta | Pt5,Pt10 | clade C |
| Phatr3_J16982 | ORTHO01HGT002832 | Bacillariophyta | Pt5 | clade C |
| Phatr3_J8538 | ORTHO01HGT004006 | *Phaeodactylum* | Pt5 | clade C |
| Phatr3_J8596 | ORTHO01HGT004006 | *Phaeodactylum* | Pt10 | clade C |
| Phatr3_Jdraft1760 | ORTHO01HGT004006 | *Phaeodactylum* | Pt5 | clade C |
| Phatr3_EG01254 | ORTHO01HGT004006 | *Phaeodactylum* | Pt1,Pt2,Pt3,Pt9,Pt4, Pt5,Pt10,Pt6,Pt7, Pt8 | all |
| Phatr3_EG01249 | ORTHO01HGT004006 | *Phaeodactylum* | Pt1,Pt2,Pt3,Pt9,Pt4, Pt5,Pt10,Pt6,Pt7, Pt8 | all |
| Phatr3_J46000 | ORTHO01HGT005106 | CCTH+SAR | Pt4,Pt8 | clade B,D |
| Phatr3_J35856 | ORTHO01HGT005106 | CCTH+SAR | Pt4 | clade B |
| Phatr3_Jdraft1549 | ORTHO01HGT005112 | CCTH+SAR | Pt5,Pt10,Pt6,Pt7, Pt8 | clade C,D |
| Phatr3_J46957 | ORTHO01HGT005294 | CCTH+SAR | Pt5,Pt10,Pt6,Pt7, Pt8 | clade C,D |
| Phatr3_Jdraft1304 | ORTHO01HGT005514 | CCTH+SAR | Pt4 | clade B |
| Phatr3_J5780 | ORTHO01HGT005959 | CCTH+SAR | Pt6,Pt7,Pt8 | clade D |
| Phatr3_Jdraft1129 | ORTHO01HGT006980 | Ochrophytes | Pt1,Pt2,Pt3,Pt9,Pt4, Pt5,Pt10,Pt6,Pt7, Pt8 | all |
| Phatr3_Jdraft1216 | ORTHO01HGT008050 | Bacillariophyta | Pt1,Pt2,Pt3,Pt9,Pt4, Pt5,Pt10,Pt6,Pt7, Pt8 | all |
| Phatr3_Jdraft1251 | ORTHO01HGT008050 | Bacillariophyta | Pt1,Pt2,Pt3,Pt9,Pt4, Pt5,Pt10,Pt6,Pt7, Pt8 | all |
| Phatr3_Jdraft1618 | ORTHO01HGT011254 | CCTH+SAR | Pt4,Pt5,Pt10,Pt6, Pt7,Pt8 | clade B,C,D |
| Phatr3_J47785 | ORTHO01HGT104904 | *Phaeodactylum* | Pt1 | clade A |
| Phatr3_EG02620 | ORTHO01HGT105075 | *Phaeodactylum* | Pt4 | clade B |

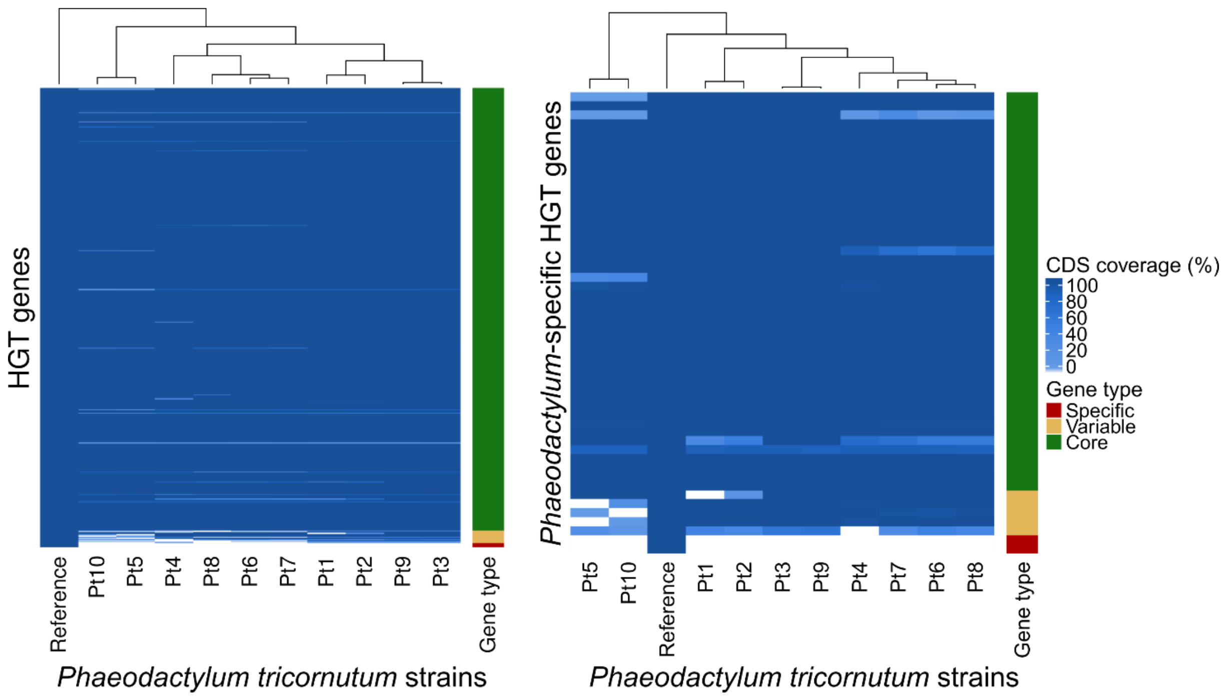

**Figure SN2: CDS coverage of HGT genes in 10 resequenced P. tricornutum strains.** CDS coverage for all (left) or species-specific (right) HGT genes across ten resequencing strains.
